# Structural and Functional Principles of Hcp-Mediated Antibacterial Toxin Delivery by the Type VI Secretion System

**DOI:** 10.64898/2026.06.15.732508

**Authors:** Po-Yin Chen, Yung-Chih Chen, Chun-Hsiung Wang, Jia-Ying Su, Chien-Ling Lin, See-Yeun Ting

## Abstract

The type VI secretion system (T6SS) is a bacterial contractile nanomachine that translocates toxic effectors into neighboring cells, helping bacteria to survive and compete in polymicrobial environments. The core T6SS structural protein Hcp forms the inner tube and mediates the loading and delivery of diverse effectors. Yet, how Hcp selectively recruits structurally and functionally distinct effectors in the absence of recognizable secretion signals remains poorly understood. Here, using *Pseudomonas aeruginosa* H1-T6SS as a model, we combined competition-coupled deep mutational scanning with cryo-EM based structural analysis to determine how Hcp accommodates distinct effectors. The resulting mutational landscape distinguishes structurally constrained residues required for Hcp tube assembly from lumen-facing residues specialized for cargo engagement. Near-atomic-resolution structures of Hcp bound to Tse2 and Tse4, together with a high-confidence Hcp–Tse1 model, revealed that while effector-specific contacts exist, Hcp engages different cargos through largely shared, dispersed interfaces characterized by polar-biased and variation-tolerant interactions. Comparative structural analyses further showed that effector-interacting residues are not governed by conserved primary-sequence motifs, but by conserved lumen-facing geometry and permissive physicochemical properties across distantly related bacteria. Together, our findings support a model in which Hcp acts as a selective yet adaptable scaffold that accommodates diverse antibacterial effectors through conserved structural features and flexible interaction surfaces, providing a mechanistic framework for understanding cargo selection by the T6SS and explaining how a conserved secretion nanomachine can evolve to deliver diverse toxic payloads.

## Introduction

Bacteria inhabit densely populated environments where competition is intense. To survive under these conditions, they have evolved diverse molecular weapons that damage neighboring cells [1]. Among these, the type VI secretion system (T6SS) is a contractile nanomachine widespread among Gram-negative bacteria that delivers diverse toxic effector proteins directly into adjacent cells through a contact-dependent manner [2–4]. An active T6SS confers a competitive advantage on bacteria within the same niche, enhancing their fitness in nutrient-limited or space-constrained environments, and influencing the compositions and dynamics of microbial communities [5–8].

The T6SS is assembled into a phage tail-like nanomachine composed of a membrane complex [9, 10], a baseplate [11], a contractile sheath [12], and an inner tube [13] capped by a spike complex [14, 15]. Upon activation, rapid contraction of the sheath propels the inner tube and spike, together with their associated effectors, through the membrane complex and into the target cell [13]. Despite progress in defining the architecture and dynamics of the T6SS, key mechanistic gaps remain in how effectors are recruited and loaded onto the molecular machine. Current models position effector loading on the secreted “puncturing device,” comprising the spike complex formed by PAAR and VgrG and the inner tube composed of haemolysin co-regulated protein (Hcp) [16]. Some T6SS effectors are fused to either VgrG, PAAR, or Hcp proteins. In other cases, some effectors, also referred to as cargo effectors, are recruited to the T6SS through protein-protein interactions with these structural components [16]. While spike-associated cargo effector loading has been characterized in considerable structural and mechanistic detail [17–20], how cargo effectors are carried by Hcp remains poorly understood. Given that Hcp forms the structural scaffold for the delivery of multiple, structurally diverse cargos during a single firing event, understanding how Hcp mediates cargo recognition is central to explaining how the T6SS achieves both specificity and payload diversity.

Hcp is a core component of the T6SS, assembling into a hexameric ring with an inner diameter of ∼40 Å [3, 21, 22], consistent with a lumen capable of accommodating cargo effectors. Stacking of these rings forms the inner tube of the contractile nanomachine, generating a repetitive array of potential effector-loading sites and enabling high-capacity delivery of an effector cocktail during a single firing event (**Extended Data Fig. 1A**). Upon T6SS firing, the Hcp inner tube—together with its loaded cargo effectors—is propelled into neighboring cells. Current models propose that the inner tube dissociates after entry into the target cell, potentially in response to changes in the intracellular environment, thereby releasing the associated effectors to exert their toxic activities. Beyond serving as a structural conduit for effector delivery, Hcp has also been shown to protect effectors from degradation through a chaperone-like activity [23, 24]. This property may ensure that effectors remain stable and readily available prior to T6SS firing. By stabilizing effectors, Hcp may buffer against premature toxicity and reduce the need for continuous effector synthesis [23, 24].

The ability of Hcp to accommodate distinct effectors, both mechanistically and spatially, underscores its role as a central hub for effector delivery. Three distinct T6SS gene clusters have been identified in the well-established T6SS model organism *Pseudomonas aeruginosa* PAO1[25]. Among them, the H1-T6SS plays a dominant role in interbacterial competition and is associated with at least eight effector proteins[26]. Four of these effectors — Tse1, Tse2, Tse3, and Tse4 — have been reported to interact with Hcp1 (PA0085, the Hcp of H1-T6SS) [23]. Interestingly, the Hcp-associated effectors exhibit distinct protein structures, biochemical activities, and cellular targets (**Extended Data Fig. 1A**). Tse1, Tse3, and Tse4 act in the periplasm, where they hydrolyze peptidoglycan cross-links[27], cleave the glycan backbone of peptidoglycan [27], and induce membrane depolarization and ion imbalance[28, 29], respectively. In contrast to these cell envelope–targeting enzymes, Tse2 is a cytoplasmic toxin proposed to disrupt essential intracellular processes, resulting in bacteriostasis or growth arrest rather than immediate cell lysis [2, 30, 31]. Although previous studies have suggested that several Hcp1 residues may be involved in effector binding, how these interactions are biased or coordinated at the molecular level remains unresolved. In particular, it is unclear how many residues within Hcp1 are functionally engaged in effector delivery, whether these residues form discrete interaction interfaces or dispersed permissive surfaces, and how they are organized within the Hcp1 structure. Moreover, though direct Hcp1-effector interactions have been demonstrated previously [23, 24], the functional contributions of individual Hcp1 residues to effector delivery have not been systematically evaluated in the context of interbacterial competition. Consequently, our understanding of how Hcp1 supports effector loading remains fragmented, largely due to the lack of unbiased, residue-resolved approaches that link molecular interactions to functional outcomes.

Here, we present a comprehensive approach to systematically assess the functional impact of mutational perturbation of Hcp1 within its native interaction network in *P. aeruginosa* PAO1. By integrating this functional landscape with cryo-EM-based structural analyses of Hcp1 in complex with its cargo effectors, we identify shared interfaces that mediate effector binding. We further extend this analysis to Hcp-effector pairs from diverse bacterial species to uncover conserved features underlying cargo engagement. Taken together, these results provide a mechanistic framework for understanding how Hcp functions as a selective scaffold for effector loading, revealing a general interaction mode governing Hcp-mediated cargo delivery across diverse T6SSs.

## Results

### A high-throughput functional profiling of hcp1 during interbacterial competition

To gain a more comprehensive understanding of the complex interactions between Hcp1 and its cargo effectors, we decided to systematically identify the important residues involved. Deep mutational scanning (DMS) enables unbiased, residue-level functional mapping by quantifying the effects of global mutational perturbations in a high-throughput manner [32]. Therefore, we applied DMS to identify residues required for Hcp1’s function in effector interaction and delivery. The T6SS plays a critical defensive role when bacteria are themselves targeted by neighboring cells. *P. aeruginosa* PAO1 has been shown to mount a rapid T6SS-mediated counterattack upon sensing assaults from competing bacteria [33]. For example, *Enterobacter cloacae* actively antagonizes PAO1 through its own T6SS, and wild-type PAO1 survives this encounter significantly better than a strain lacking *hcp1* [34] (**Extended Data Fig. 1B**). Leveraging Hcp1’s role in this defensive counterattack, we established a high-throughput functional screen based on interbacterial competition between PAO1 and *E. cloacae* (**Fig. 1A**). During co-incubation, attacks from *E. cloacae* trigger a retaliatory T6SS response in PAO1, such that the survival of each Hcp1 variant reflects its ability to support effective T6SS function, including efficient effector delivery (**Fig. 1B**). Consistent with this interpretation, a PAO1 strain lacking all four reported Hcp1-associated cargo effector genes (*tse1–4*, referred to as Δ*cargos*) exhibited significantly reduced survival compared with the wild type, yet higher survival than the Δ*hcp1* strain, indicative of partial T6SS dysfunction (**Fig. 1C**). These observations support interbacterial competition as a potentially quantitative readout for assessing Hcp1 function during T6SS-mediated antagonism.

**Fig. 1:**
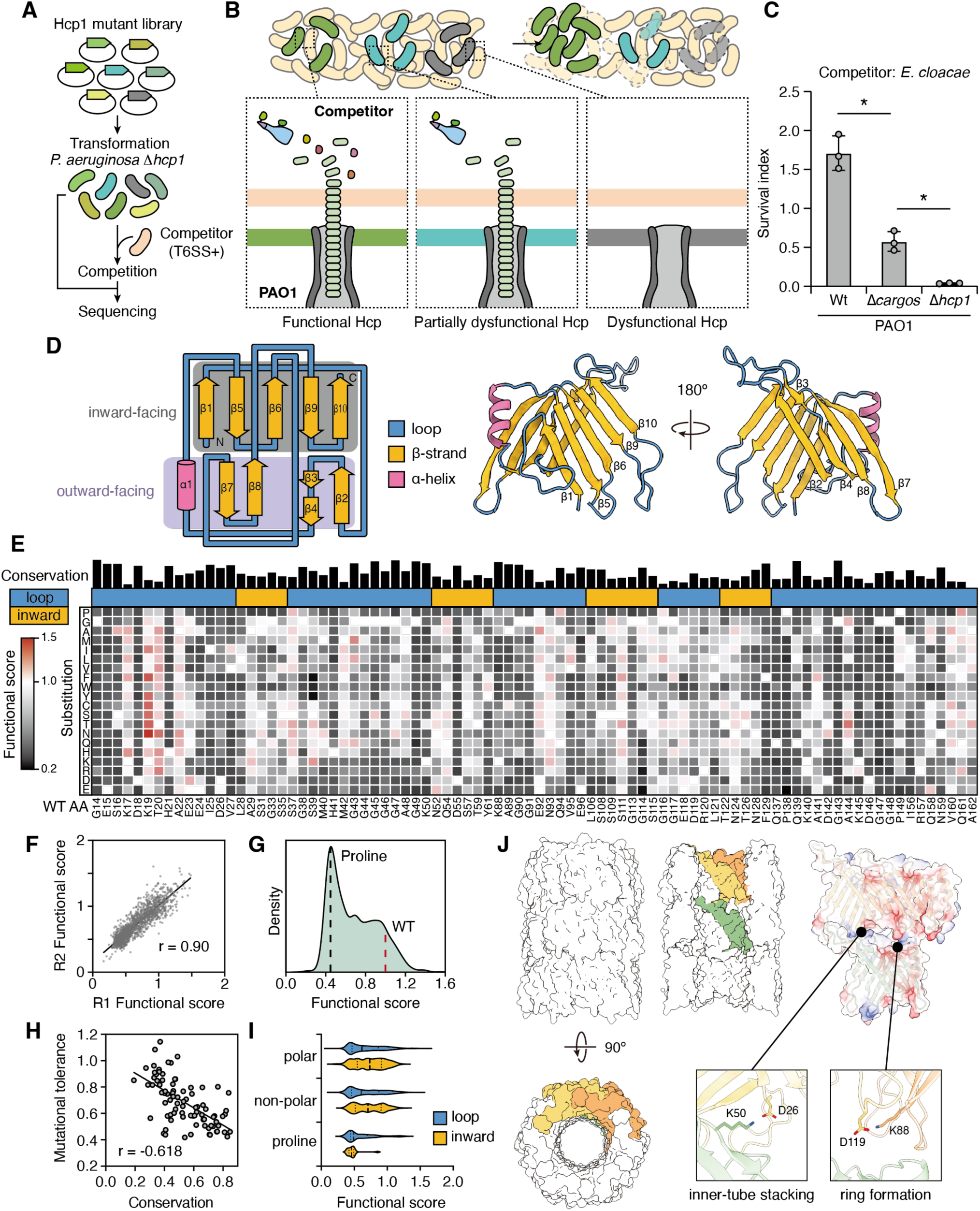
A deep mutational scan individually characterizes single-amino acid mutations in Hcp1 during interbacterial competition. (A) Schematic of the DMS workflow. A library of Hcp1 single-amino acid substitution variants was transformed into PAO1 Δ*hcp1* and subjected to interbacterial competition against a T6SS-active competitor, *E. cloacae*. Variant frequencies were determined by sequencing before and after competition. Functional scores were calculated as the relative survival of each Hcp1 variant following competition, normalized to wild-type Hcp1. (B) Schematic illustrating PAO1 strains harboring functional, partially dysfunctional, or dysfunctional Hcp1 variants during interbacterial competition. In this context, functional Hcp1 supports both T6SS assembly and cargo effector delivery, resulting in higher survival. Partially dysfunctional Hcp1 variants retain T6SS assembly and firing but exhibit impaired cargo loading or delivery, providing intermediate survival. Dysfunctional Hcp1 variants fail to support inner-tube assembly and consequently abolish effector delivery, leading to the lowest survival. (C) Survival of PAO1 strains following 6-hour co-culture with *E. cloacae*. Data are presented as mean ± SD. n = 3 biological replicates with three technical replicates each. Asterisks indicate statistically significant differences (p < 0.05, two-tailed t-test). (D) Topology and three-dimensional structure of Hcp1 (PDB = 1Y12). The grey area represents the inward-facing region; the purple area represents the outward-facing region. (E) Heatmap of the Hcp1 deep mutational scanning dataset. Columns represent residue positions in Hcp1 and rows represent amino acid substitutions. Colors indicate functional scores derived from interbacterial competition assays, with higher scores shown in red and lower scores shown in black. Sequence conservation at each position is displayed above the heatmap, and residue locations within the lumen-facing (inward) surface or loop regions of the Hcp1 ring are indicated by the corresponding annotations. (F) Correlation between two representative biological replicates. Each point represents a single variant. See also Extended Data Fig. 1C. R1 and R2: replicate 1 and 2. (G) Density distribution of functional scores for all variants. The red dashed line indicates the wild-type score (1.0). The black dashed line indicates the mean score of proline substitutions located within β-strand regions. (H) Relationship between positional conservation and mutational tolerance. Mutational tolerance was calculated as the mean functional score of all substitutions at each position. The black line represents a linear regression fit. (I) Violin plots depicting the distribution of functional scores of variants carrying substitutions to polar, nonpolar, or proline residues. The width of each plot is proportional to the density of data points for this range of values, and the horizontal stripes mark the quartiles. (J) The Hcp1 tubular model (see also Extended Data Fig. 2). Three representative subunits are highlighted in green, yellow and orange. Residues involved in hexamer formation (K88 and D119) and inter-ring stacking (D26 and K50) are indicated.

Next, we generated a comprehensive Hcp1 variant library comprising 1,634 variants, representing all 19 non-synonymous substitutions at 86 Hcp1 residues. Residues with side chains facing the exterior of the Hcp1 hexamer were excluded from library design to minimize potential effects on Hcp1-sheath interactions during T6SS assembly [35] (**Fig. 1D**). Plasmids encoding the Hcp1 variants were introduced into a *P. aeruginosa* PAO1 Δ*hcp1* strain to assess mutant fitness. Sequencing of the pre-competition population recovered all designed variants across the three biological replicates, with median coverages exceeding 1,100 reads per variant, indicating complete and highly uniform representation of the library prior to competition.

After competition, functional scores for each Hcp1 variant were calculated by normalizing pre- and post-selection frequency ratios against wild-type Hcp1 (**Fig. 1E**), generating a residue-level functional landscape of Hcp1. Our screen was highly reproducible across three biological replicates, with pairwise Pearson correlation coefficients ranging from 0.81 to 0.90 (**Fig. 1F and Extended Data Fig. 1C**). Functional scores ranged from 0.17 to 1.55 (mean = 0.68; median = 0.63), with 72.7% of the variants displaying reduced fitness relative to wild type. The distribution was markedly asymmetric, featuring a dominant peak at ∼0.4–0.5 and a long rightward tail toward higher scores (**Fig. 1G**), indicating that most substitutions impair Hcp1 function, with a small subset being better tolerated. The non-normal distribution and the presence of a secondary shoulder in the mid-range of functional scores further evidence heterogeneous mutational sensitivities across positions. Notably, proline substitutions within β-strand regions displayed an average score of 0.47, closely aligning with the principal peak of the distribution. Given the well-established disruptive effect of proline on secondary structure, this pattern suggests that backbone perturbation consistently imposes substantial functional constraints across Hcp1. As expected, average score at each position negatively correlated with sequence conservation (Pearson’s r = -0.62) (**Fig. 1H**), indicating that the mutational landscape captured by our screen reflects evolutionary constraints acting on Hcp1.

Interestingly, our functional screen revealed that certain regions of the Hcp1 structure are more sensitive to specific types of mutations than others. For example, loop regions are more sensitive to mutation than residues with inward-facing side chains (**Fig. 1I**), highlighting the structural constraints required for Hcp1 ring stacking [21, 36]. Notably, residues D26 and K50—predicted to form key electrostatic interactions for ring stacking—were highly sensitive to most substitutions; yet, they remained relatively permissive to mutations that preserved the original charge (**Fig. 1J, Extended Data Fig. 2**). Similarly, K88 and D119, located on adjacent Hcp1 subunits, appear to facilitate ring formation via electrostatic attraction (**Fig. 1J, Extended Data Fig. 2**), as evidenced by the low functional scores associated with most substitutions and the relatively high tolerance for mutations that preserved the original charge. Collectively, these results confirm that the DMS dataset is biologically robust, capturing critical evolutionary and structural constraints essential for Hcp1 function during interbacterial competition.

### Unsupervised learning reveals functionally distinct classes of Hcp1 residues

Our DMS dataset provides a comprehensive, residue-level view of Hcp1’s functional response to amino acid substitutions. To better understand groups of residues with similar mutational response profiles that may reflect shared structural or biochemical constraints in effector delivery, we applied unsupervised learning to the Hcp1 mutational landscape. [37, 38]. Specifically, for each residue, substitution profiles incorporating physicochemical features and functional scores were embedded using Uniform Manifold Approximation and Projection (UMAP) [39], before being subjected to density-based hierarchical clustering (HDBSCAN) [40], which resolved six distinct mutational response clusters (**Fig. 2A, Supplementary Table 1**).

**Fig. 2:**
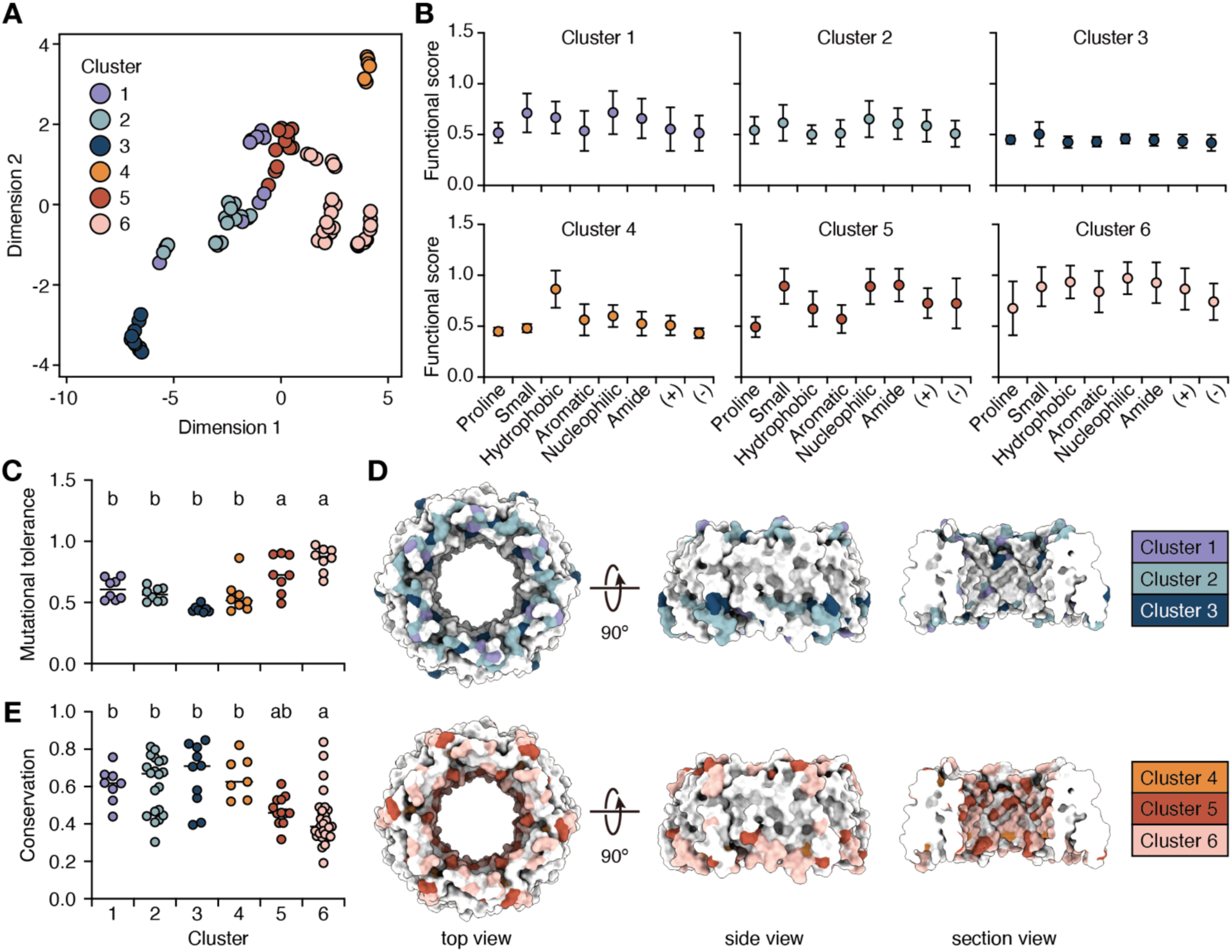
Unsupervised learning identifies residue clusters with distinct mutational response profiles. (A) UMAP projection of Hcp1 residues based on their mutational response profiles. Each point represents a residue, and colors indicate the six clusters defined by HDBSCAN. (B) Mean functional scores of residues within each cluster following substitutions belonging to different physicochemical classes. (C) Mutational tolerance of residues within each cluster. (D) Mapping of cluster assignments onto the Hcp1 hexamer structure (PDB: 1Y12). (E) Sequence conservation of residues within each cluster. Different letters above groups indicate differences among clusters (one-way ANOVA followed by Tukey’s multiple comparisons test, p < 0.05).

To define the characteristic features of each cluster, we quantified the average functional scores associated with different classes of amino acid substitutions (**Fig. 2B**). This analysis revealed three principal modes of mutational sensitivity. First, residues in Clusters 1–3 exhibited uniformly low functional scores (mean 0.44–0.62) across most substitutions, indicating broad intolerance to amino acid changes (**Fig. 2C**). Second, two clusters displayed clear biochemical preferences: residues in Cluster 4 preferentially tolerated hydrophobic substitutions, whereas those in Cluster 5 favored polar substitutions (**Fig. 2B**), indicative of chemically specific interactions. Finally, residues in Cluster 6 were broadly tolerant but showed pronounced sensitivity to proline, consistent with structural constraints imposed by local backbone geometry.

Next, we projected the mutational response clusters onto the Hcp1 ring structure, revealing a pronounced non-random spatial organization (**Fig. 2D**). Globally intolerant residues (Clusters 1–3) were enriched at the top and bottom of the ring, representing regions critical for inter-ring stacking and structural stability. Consistently, these residues proved highly conserved across Hcp homologs, supporting their role in maintaining structural integrity during T6SS assembly (**Fig. 2E**). In contrast, chemically selective and broadly tolerant residues (Clusters 4–6) were preferentially localized in the inner channel, implying functional specialization beyond core assembly. Notably, residues in Clusters 5 and 6 exhibited lower sequence conservation across diverse Gram-negative bacteria, consistent with the idea that lumen-facing regions can accommodate sequence diversification without compromising function (**Fig. 2E**). Importantly, the assignment of these residues to lumen-facing clusters was not predefined but emerged from their shared mutational signatures, indicating that their functional properties, rather than prior structural assumptions, underlie this segregation. The combination of lumen-facing localization, selective mutational tolerance, and reduced sequence conservation is characteristic of adaptable protein–protein interaction interfaces, pointing to Clusters 5 and 6 being involved in effector engagement rather than core structural stability. Consistent with this interpretation, we noted that previously identified secretion-relevant residues mapped to these clusters: S31, which selectively contributes to Tse2 secretion, falls within Cluster 5; whereas T59 and S115 that are required for delivery of multiple effectors (Tse1–Tse4), belong to Cluster 6 [23, 24]. Together, these results reveal a functional partitioning of the Hcp1 structure. While conserved and mutationally intolerant residues are concentrated at regions required for ring assembly and tube stacking, chemically selective and mutationally permissive residues define a distinct lumen-facing surface enriched for cargo-interacting determinants. This organization suggests that Hcp1 maintains a robust structural scaffold while preserving an adaptable interface capable of accommodating diverse cargo effectors.

### Cryo-EM of the Hcp1–Tse2 complex reveals the structural basis of Hcp1-mediated cargo engagement

Although our UMAP analysis revealed distinct mutational response patterns across the Hcp1 functional landscape, it could not directly pinpoint the residues responsible for effector interaction. To overcome this limitation, we reasoned that structural information may provide a complementary framework for identifying effector-contacting residues and anchoring functional patterns observed in the global mutational profile. To pursue this avenue, we determined the structure of Hcp1 in complex with Tse2, a well-characterized cargo effector whose stabilization and secretion are strictly Hcp1-dependent, making it a robust model for defining cargo–Hcp interaction interfaces [24].

The Hcp1–Tse2 complex was heterologously expressed and purified from *E. coli* and analyzed by single-particle cryogenic electron microscopy (cryo-EM), yielding a reconstruction at an average resolution of 2.73 Å (**Fig. 3A and Extended Data Figs. 3 and 4**). The structure reveals a canonical Hcp1 hexameric ring forming a central lumen, consistent with the previously resolved crystal structure (PDB: 1Y12) [3]. Comparison of Hcp1 rings in the presence and absence of Tse2 shows minimal global rearrangement upon cargo loading (Cα RMSD = 0.753 Å), indicating that Tse2 binding does not induce large-scale alterations in ring architecture (**Extended Data Fig. 4**).

**Fig. 3:**
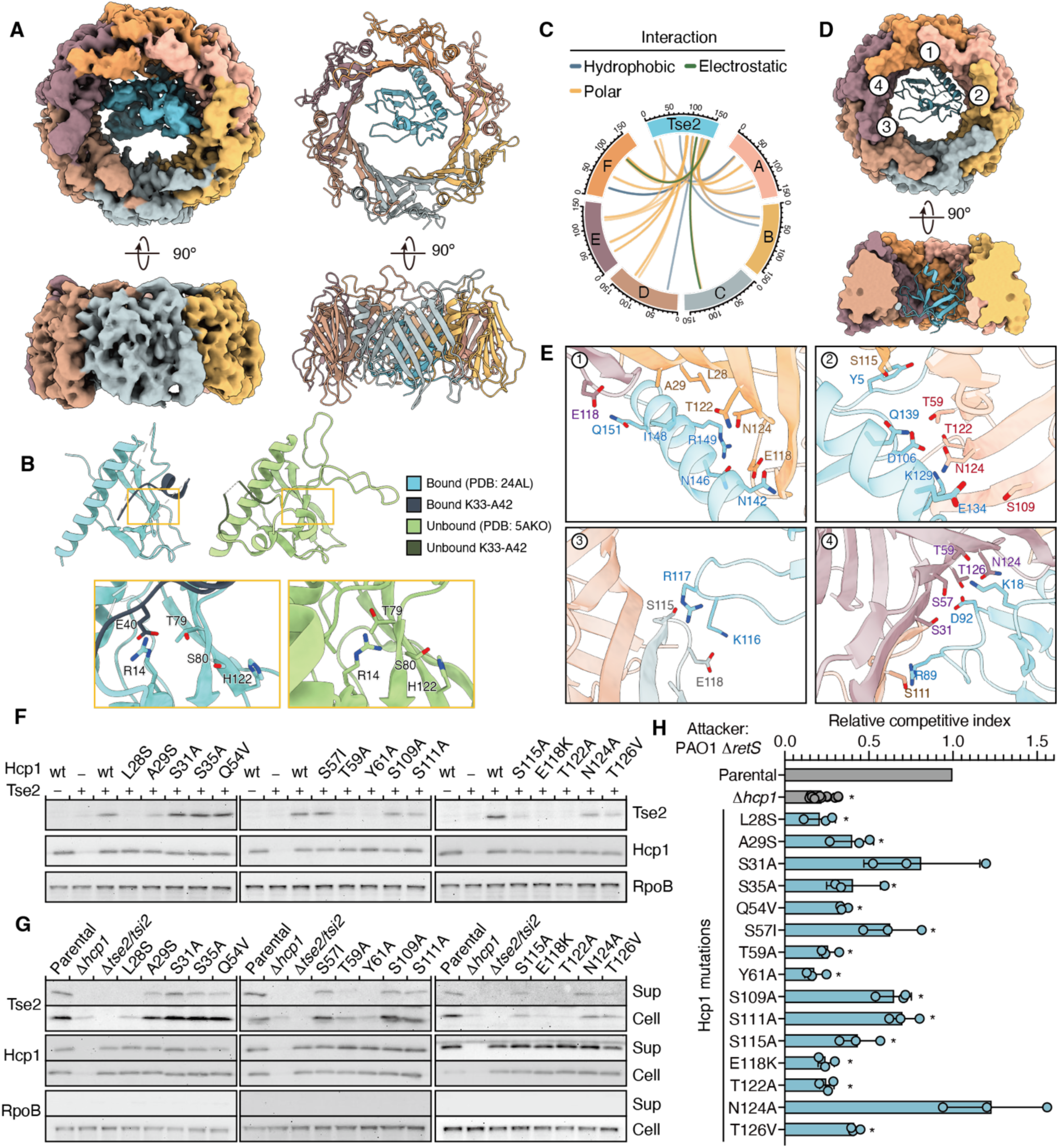
Cryo-EM analysis and functional validation of Hcp1-Tse2 complex. (A) Cryo-EM density map (left) and the corresponding atomic model (right) of Hcp1-Tse2 complex (PDB: 24AL). Dash lines represent the unresolved region lacking density. (B) Tse2 structure in Hcp1-bound (blue) and unbound forms (green). Yellow boxes highlight the catalytic pocket of Tse2 (R14, T79, S80, and H122) and the region undergoing conformational rearrangement upon Hcp1 binding. (C) Circular visualization of the structural interaction landscape between the six Hcp1 ring subunits and Tse2. Outer rings represent individual protein sequences with residue positions indicated by axis ticks. Internal links represent specific atomic interactions, color-coded by interaction types: polar (yellow), hydrophobic (blue), and electrostatic (green) interactions. (D) Atomic model in cartoon representation with numbers corresponding to the interfaces shown in detail in (E). (E) Interfaces of the Hcp1-Tse2 interaction. (F) Tse2 stability assay. Each hcp1 variant was co-expressed with a catalytic inactive Tse2 (T79A/S80A) in *E. coli*. Immunoblot results showing the effect of each Hcp1 point mutation on intracellular levels of Tse2. (G) Tse2 secretion assay of PAO1 strains having *hcp1* mutations. Immunoblot results showing the effect of each Hcp1 point mutation on intracellular and extracellular levels of Tse2 and Hcp1. The loading control is RNA Polymerase β protein (RpoB). Sup: supernatant; Cell: bacterial cell pellet. (H) Relative competitive index of PAO1 attacker strains having *hcp1* mutations. PAO1 strain with *tse2*/*tsi2* gene deletion was used as the recipient. Cells were cocultured at a 10:1 attacker-to-recipient ratio for 6 hours. Data are presented as means ± SD. n = 3 biological replicates with three technical replicates each. Asterisks indicate statistically significant differences between the indicated mean values (p < 0.05, two-tailed t-test).

**Fig. 4:**
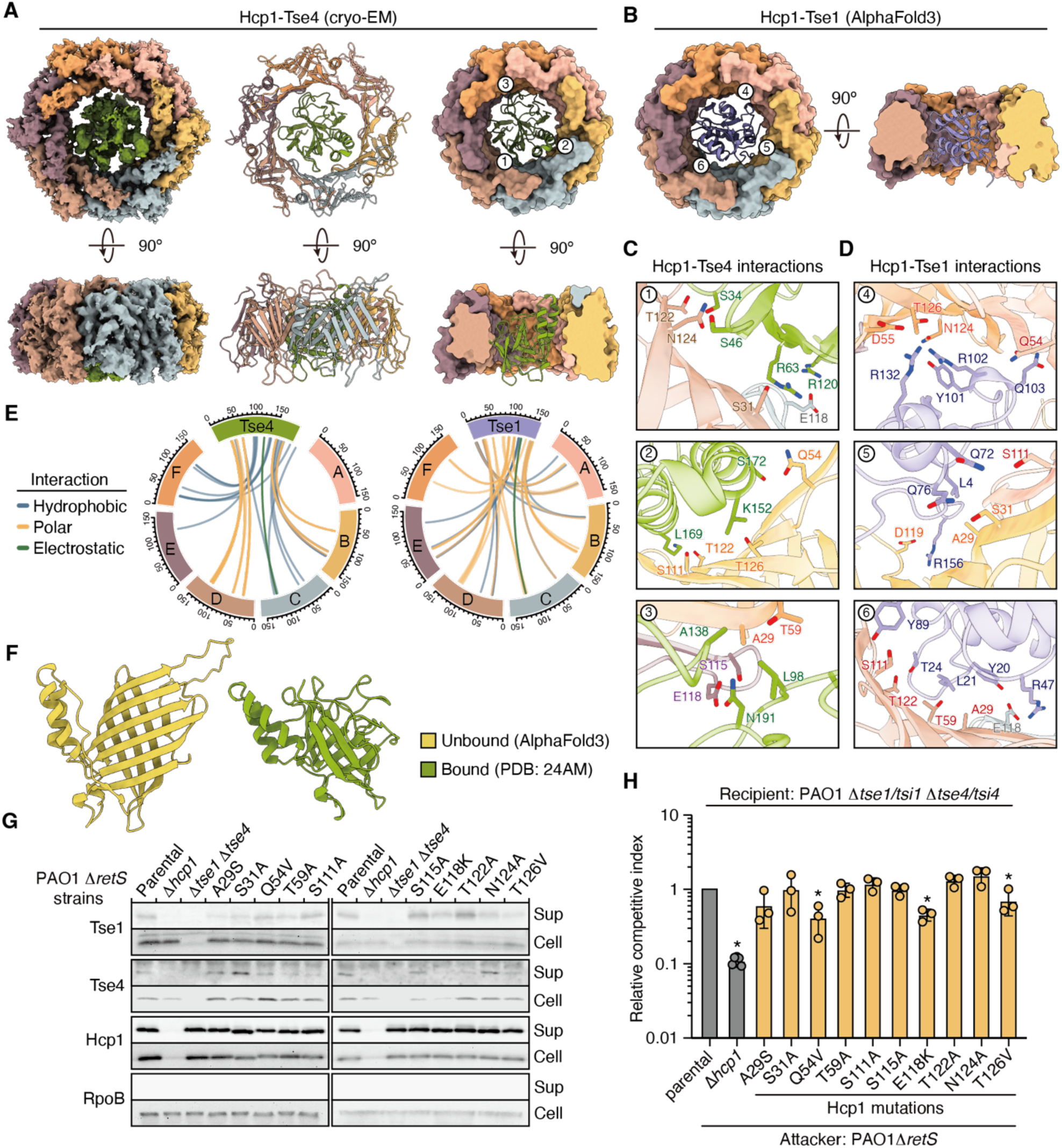
Structural analysis on Hcp1 in complex with Tse1 or Tse4. (A) Cryo-EM density maps (left) of Hcp1-Tse4 complex and the corresponding atomic models (middle and right). Numbers corresponding to the interfaces shown in detail in (C). (B) Alphafold3-predicted model of Hcp1-Tse1. Numbers corresponding to the interfaces shown in detail in (D). (C-D) Interfaces of the Hcp1-Tse4 interaction (C) and Hcp1-Tse1 (D) interaction. (E) Circular visualization of the structural interaction landscape between the six Hcp1 ring subunits and Tse4 (left) or Tse1 (right). Outer rings represent individual protein sequences with residue positions indicated by axis ticks. Internal links represent specific atomic interactions, color-coded by interaction types: polar (yellow), hydrophobic (blue), and electrostatic (green) interactions. (F) Structure of Tse4 when bound to Hcp1 ring (green) and not bound to the ring (yellow). (G) Immunoblot results showing the effect of each Hcp1 point mutation on intracellular and extracellular levels of Tse1, Tse4, and Hcp1. The loading control is RNA Polymerase β protein (RpoB). Sup: supernatant; Cell: bacterial cell pellet. (H) Relative competitive index of PAO1 attacker strains having *hcp1* mutations. Cells were cocultured at a 10:1 attacker-to-recipient ratio for 6 hours. Data are presented as means ± SD. n = 3 biological replicates with three technical replicates each. Asterisks indicate statistically significant differences between the indicated mean values (p < 0.05, two-tailed t-test).

Interestingly, structural comparison of Tse2 in its Hcp1-bound form with the previously resolved unbound crystal structure (PDB: 5AKO) revealed a localized conformational change (**Fig. 3B**). In the Hcp1-bound state, residues K33–A42 of Tse2 adopt a short α-helix followed by a loop that folds inward toward the protein core, resulting in a more “closed” conformation. However, in the unbound structure, this region extends outward and is more solvent-exposed. We also noticed that, in the Hcp1-bound form, the K33–A42 segment of Tse2 is positioned proximal to its predicted catalytic pocket, formed by amino acids R14, T79, S80, and H122 [30, 31]. This structural rearrangement supports that cargo loading into the Hcp1 lumen may induce a conformational state in Tse2 that partially occludes its active site, potentially suppressing toxicity prior to secretion. Consistent with this model, residue E40 of Tse2 lies in close proximity to R14 in our Hcp1-Tse2 complex structure (**Fig. 3B**), raising the possibility of an electrostatic interaction that stabilizes this inactive conformation.

Tse2 is positioned asymmetrically within the Hcp1 lumen, with its α-helix spanning residues R133–R156 forming the primary interface with two adjacent Hcp1 subunits (**Figs. 3C-3E**). Structural analysis identified 15 Hcp1 residues (i.e., L28, A29, S31, S35, Q54, S57, T59, Y61, S109, S111, S115, E118, T122, N124, and T126) that make direct contacts with Tse2. The Hcp1-Tse2 interface is dominated by polar interactions, with additional hydrophobic and electrostatic contributions. This composition implies a binding surface that favors reversible contacts, consistent with the requirement for stable cargo loading prior to secretion while permitting release upon T6SS firing.

### Distinct Hcp1 interface residues differentially contribute to Tse2 stabilization, secretion, and delivery

To evaluate the contribution of individual interface residues to Tse2 interaction, we generated single amino acid substitutions at each identified Hcp1 contact site. To limit global structural perturbations while allowing residue-specific functional assessment, hydrophobic residues were replaced with serine, whereas polar residues were substituted with alanine or valine. The effects of these substitutions on cargo interaction were first examined by monitoring Tse2 stability during co-expression with Hcp1 in *Escherichia coli* [24]. To avoid toxicity to the host cells, we introduced the catalytically inactive mutations (T79A/S80A) into Tse2 in the assay.

As expected, co-expression of Tse2 with wild-type Hcp1 markedly stabilized Tse2. However, several Hcp1 variants (L28S, A29S, T59A, Y61A, E118K, and T122A) substantially reduced Tse2 abundance despite comparable Hcp1 expression levels (**Fig. 3F**), consistent with impaired complex formation and/or decreased cargo stabilization. A subset of mutants (S109A, S111A, S115A, and T126V) caused a modest reduction in Tse2 levels, implying partial disruption of the Hcp1–Tse2 interaction. In contrast, five substitutions (S31A, S35A, Q54V, S57I, N124A) had no detectable impact on Tse2 accumulation, suggesting that their contributions to Tse2 binding or stabilization are either minimal or functionally redundant.

We wondered if, in the native system of *P. aeruginosa* PAO1, these Hcp1 residues also function similarly with respect to effector stability, secretion, and delivery. Therefore, we introduced the same substitutions into the chromosomal *hcp1* locus of a T6SS-hyperactive PAO1 strain lacking the negative regulator RetS [2]. To assess Tse2 stabilization and secretion, cell pellets and culture supernatants were collected for immunoblot analysis. Hcp1 protein levels in both the cellular and secreted fractions were comparable across all variants, indicating that these substitutions did not impair Hcp1 production or secretion. We observed that intracellular Tse2 levels varied across the different Hcp1 variants. Mutations (L28S, T59A, Y61A, E118K, T122A, and T126V) that significantly diminished Tse2 stability in cells also prevented its secretion. Conversely, substitutions S31A and N124A, which did not affect intracellular Tse2 levels, elicited secretion levels comparable to wild type (**Fig. 3G**).

We noticed that two substitutions, S35A and Q54V, did not measurably affect intracellular Tse2 stability but resulted in a pronounced reduction in Tse2 secretion. This uncoupling of cargo stabilization from secretion efficiency suggests that these residues are not strictly required for Hcp1–Tse2 binding but may instead function at a subsequent step in the delivery process. One possibility is that S35 and Q54 facilitate productive cargo release during or following T6SS firing. Alternatively, these residues may influence Tse2 positioning within the Hcp1 lumen or they modulate subtle aspects of tube assembly or dynamics that are critical for efficient secretion.

To assess the functional consequences of these mutations during interbacterial competition, the same Hcp1 variants were used as attacker strains against a Tse2-sensitive recipient lacking Tse2 and the cognate immunity protein Tsi2 [2]. Competitive outcomes closely mirrored the stability of intracellular Tse2. Deletion of *hcp1* abolished both Tse2 stability and Tse2-mediated killing. Similarly, variants with mutations that abolished Tse2 secretion exerted minimal killing activity, comparable to that of the Δ*hcp1* strain. Variants with partial reduction in Tse2 secretion showed a corresponding decrease in competitive fitness (**Fig. 3H**). In contrast, the S31A and N124A variants, which retained normal Tse2 stability and secretion, showed little or no reduction in killing efficiency. These results indicate that individual interface residues contribute unequally to cargo delivery, revealing partial functional redundancy within the Hcp1 binding surface.

### Extended structural analyses of Hcp1-Tse4 and Hcp1-Tse1 identify shared Hcp1 interaction sites across diverse effectors

From our structural analysis, we identified 15 Hcp1 residues that interact with Tse2. To verify whether these residues share any structural or biochemical features important for their participation in effector delivery during interbacterial competition, we mapped them back to the UMAP clusters defined by our DMS mutational profiles. Interestingly, 13 of the 15 Hcp1 residues contacting Tse2 fall into two of the UMAP clusters (Clusters 5 and 6) (**Fig. 2A and Extended Data Fig. 5**). This grouping supports our supposition that these clusters represent essential features for loading effectors, including their polar-tolerant chemical traits and placement within the lumen, which determine how Hcp1 recognizes cargo effectors. Next, to determine if these features extend beyond Tse2, we used structural approaches to study Hcp1’s interactions with other cargo effectors, i.e., Tse1, Tse3, and Tse4. Following the same pipeline as deployed for our Hcp1-Tse2 analysis, we successfully purified the Hcp1–Tse4 complex and resolved its structure by cryo-EM at 2.83 Å resolution (**Fig. 4A and Extended Data Figs. 6 and 7**). Unfortunately, we were unable to obtain high-quality complex structures for Hcp1–Tse1 or Hcp1–Tse3 due to technical limitations, so instead we applied AlphaFold3 models to predict the structures of these two complexes [41]. In doing so, we obtained a predicted model of Hcp1–Tse1 complex with high confidence scores (ipTM = 0.76, pTM = 0.79) (**Fig. 4B**), but confidence scores were much lower for Hcp1–Tse3, (ipTM = 0.57, pTM = 0.61). One possible explanation for this discrepancy is the larger size of Tse3’s (44.4 kDa), which may require a non-canonical mode of interaction that is more difficult for current structure-prediction methods to capture. Consequently, we focused our subsequent analyses on the interactions of Hcp1 with Tse1 and Tse4.

By analyzing the newly resolved cryo-EM structure of Hcp1–Tse4, we found that, unlike Tse2, binding of witch primarily occurs with two Hcp1 subunits, Tse4 is less asymmetrically positioned within the Hcp1 lumen, contacting five of six Hcp1 subunits. In addition to effector position within the lumen, similar to Hcp1-Tse2, the interactions between Hcp1 and Tse4 are primarily polar, i.e. similar to Hcp1-Tse2, with hydrophobic and electrostatic interactions also contributing (**Figs. 4B, 4D, and 4E**). A previous biochemical study found that Tse4 acts as a pore-forming toxin that integrates into bacterial membranes, creating ion-selective channels that cause ion leakage, leading to membrane depolarization and subsequent bacterial death [28, 29]. Here, we provide the first structural view of Tse4 bound to Hcp1. To compare the structures of Tse4 with or without binding within the Hcp1 ring, we used AlphaFold3 to obtain a predicted model of the Tse4 protein structure alone [41]. The resulting Tse4 model is comprised of an eight-stranded β-barrel followed by two α-helices (**Fig. 4F**). The β-barrel is primarily hydrophobic (**Extended Data Fig. 8A**), consistent with Tse4’s membrane-targeting capability. Surprisingly, in our Hcp1-bound structure we resolved from cryo-EM, the β-barrel appears shorter and is flanked by flexible loops (**Fig. 4F**), with the surface mostly being hydrophilic (**Extended Data Fig. 8A**). This structural discrepancy indicates that Tse4 may adopt a hydrophilic conformation within the Hcp1 lumen. We proposed that, as in Tse2, this conformational change may be required for Tse4 upon loading within the Hcp1 ring. One possibility is that this conformational change allows Tse4 to present a surface of lower hydrophobicity that can be more readily accommodated within the Hcp1 inner tube, which is highly hydrophilic (**Extended Data Fig. 8B**). Furthermore, this change in surface properties may also inhibit Tse4’s membrane-insertion activity, in which the hydrophobic surface plays a major role, before its release into the target cell. Although direct evidence for such a conformational switch is currently lacking, this model provides a conceivable framework linking Hcp-mediated cargo accommodation with the membrane-disruptive activity of Tse4.

The predicted Hcp1-Tse1 complex presents a pattern similar to that of Hcp1-Tse4, with interaction sites occurring on all six Hcp1 units. As observed for Tse2 and Tse4, most connections between Tse1 and Hcp1 are via polar bonds, with some hydrophobic and electrostatic links (**Figs. 4C-4E**). We did not observe dramatic differences between Hcp1-bound Tse1, as resolved in our AlphaFold3-predicted model, and Tse1 not bound to Hcp1 (PDB: 4FGE) (**Extended Data Fig. 8C**). As for Tse2, more than 85% of the Hcp1 residues that contact Tse1 or Tse4 are in UMAP-defined Clusters 5 and 6 (**Extended Data Fig. 5**), highlighting a common interaction pattern in which the polar interactions of Hcp1 lumen-facing residues predominate, and their tolerance to mutations is relatively high compared to the inner tube structure-related residues. Given these common features, we wondered if Hcp1 employs a set of shared residues for effector interaction. To explore this possibility, we compared our three Hcp1-effector complex structures we obtained. Interestingly, we identified 10 Hcp1 residues shared in all of the effector complexes: A29, S31, Q54, T59, S111, S115, E118, T122, N124, T126). To evaluate the roles of these shared residues in Tse1 and Tse4 secretion, we performed secretion assays as described above. We found that compared to the parental strain expressing wild-type Hcp1, the secretion of Tse4 was strongly abolished by a subset of mutations (T59A, S111A, E118K, T122A, and T126V), whereas the other mutations elicited either a minor or no effect (A29S, S31A, Q54V, S115A, N124A). The effects of these mutations on Tse1 secretion were less obvious, either prompting a slight decrease (A29S, S31A, T59A, E118K, N124A, T126V) or no effect (Q54V, S111A, S115A, T122A) (**Fig. 4G**).

In addition, unlike the intracellular protein level of Tse2, which was dramatically affected by these mutations, we observed no effect of any of the mutations on the intracellular protein level of Tse1, and only the E118K substitution had a slightly negative impact on the protein level of Tse4. This outcome indicates that the degree of each effector’s reliance on Hcp1’s chaperone activity varies.

We further assessed the contribution of Hcp1 residues to effector delivery during interbacterial competition. However, robust killing phenotypes are not always observed for individual effectors. Consistent with previous reports, deletion of the cognate immunity protein against either Tse1 or Tse4 (Tsi1 and Tsi4, respectively) alone did not produce a strong killing phenotype [27, 28], indicating that neither effector is sufficient on its own to generate a robust killing readout. To overcome this limitation, we adopted an alternative strategy based on prior evidence that Tse1 and Tse4 act synergistically to enhance bacterial killing [28]. Accordingly, instead of using recipients lacking a single immunity gene, we employed a recipient strain deficient in both Tsi1 and Tsi4. This sensitized background enabled detection of the combined activity of the two effectors and provided a robust phenotype for evaluating the impact of Hcp1 interface mutations on effector delivery (**Extended Data Fig. 8D**). Using this assay, two of the ten Hcp1 variants tested (E118K and T126V) exhibited reduced competitiveness relative to the parental strain, consistent with impaired delivery of both Tse1 and Tse4. In contrast, variants that selectively affected the secretion of a single effector (A29S, S31A, T59A, S111A, S115A, T122A, and N124A) did not exhibit significant competitive defects. Interestingly, the Q54V mutant displayed reduced competitiveness despite largely preserved Tse1 secretion and only partial reduction of Tse4 secretion. One possible explanation for this outcome is that this mutation affects a later step of effector delivery or release. For example, this mutation may increase Hcp1 binding affinity to effectors, thereby making effector release harder when translocated into the target cells (**Fig. 4H**). Together, by combining high-resolution structures of Hcp1 and cognate effectors, along with mutational studies, our data delineate a conserved effector-loading mode, whereby Hcp1 utilizes a specific set of residues to accommodate diverse cargo effectors, including Tse1, Tse2, and Tse4.

### Structure-guided and sequence-permissive principles of Hcp1–cargo recognition

Among *P. aeruginosa* isolates, Hcp1 is highly conserved at the amino acid level, including the effector-interacting residues we identified through deep mutational scanning (DMS) and structural analyses (**Extended Data Fig. 9A**). This high conservation implies a shared intra-species strategy for cargo loading. In contrast, our analysis of Hcp1 homologs from phylogenetically diverse bacteria revealed that while residues required for hexameric ring formation or inner-tube stacking are broadly conserved, the effector-interacting sites vary substantially across different species (**Fig. 5A**). This divergent pattern indicates that Hcp–cargo interactions are not dictated by rigid primary sequence motifs. Therefore, we hypothesized that Hcp1 utilizes a generalized structural framework, rather than sequence-specific hotspots, to recognize cargo effectors.

**Fig. 5:**
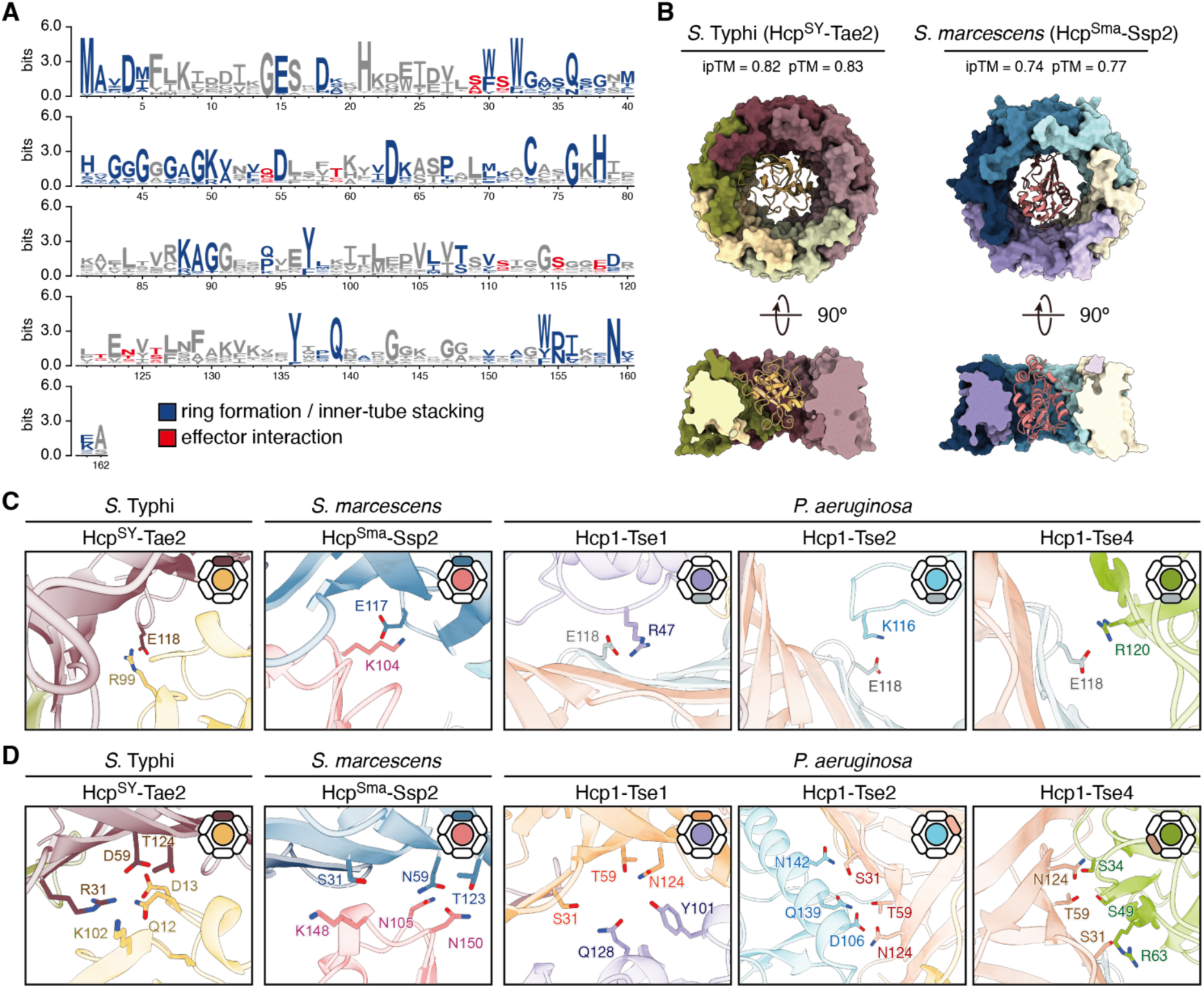
Sequence and Structural Comparison on Hcp1 homologs. (A) Sequence Logos of the Hcp1 multiple sequence alignments across homologs from phylogenetically diverse bacterial species for 1,809 protein sequences, with residues involve in inner-tube structure assembly or effector interaction colored in blue and red, respectively. (B) AlphaFold3 predicted models of Hcp^SY^-Tae2 from *S.* Typhi (left panel) and Hcp^SY^-Ssp2 from *S. marcescens* (right panel). (C-D) Conserved electrostatic interaction (C) and polar binding pocket (D) found in Hcp-effector complexes from *S.* Typhi, *S. marcescens*, and *P. aeruginosa.* The interfaces are labeled, and the corresponding Hcp1 subunits are highlighted in the ring schematics.

To test this hypothesis and understand how Hcp1 family members in other species engage their payloads, we analyzed AlphaFold3 models of two previously characterized, distantly related Hcp1 homolog–effector pairs: *Salmonella* Typhi Hcp^SY^–Tae2 and *Serratia marcescens* Hcp^Sma^–Ssp2 (**Fig. 5B**) [24, 42]. The predicted Hcp rings exhibit high structural similarity to Hcp1 of PAO1. This structural conservation is further supported by the close alignment between PAO1 Hcp1 and an experimentally resolved Hcp structure from *S.* Typhimurium (PDB: 5XHH; Cα RMSD = 0.718 Å), which shares high sequence identity with *S.* Typhi (**Extended Data Fig. 9B**). Despite these Hcp1 homologs sharing limited primary sequence identity (<50%) with PAO1 Hcp1, residues occupying equivalent lumen-facing positions in both models directly contact their cognate effectors (**Extended Data Fig. 9C**). Strikingly, we found a substantial proportion of the predicted interface residues in Hcp^SY^ (80%) and Hcp^Sma^ (75%) map to positions corresponding to the PAO1 Hcp1 interaction sites and are enriched within the same UMAP-defined clusters (Clusters 5 and 6) (**Supplementary Table 2**). These findings suport that Hcp-mediated cargo recognition is primarily governed by a conserved geometry and permissive physicochemical properties rather than strict primary sequence conservation.

Within this structural framework, we noted that several interaction features recurred across systems. In particular, we observed electrostatic interactions involving the conserved Hcp residue E118 across all examined complexes (**Fig. 5C**), suggesting that electrostatic anchoring represents a universal principle of cargo engagement. Consistent with this interpretation, mutating E118 significantly impaired the secretion and interbacterial delivery of all examined effectors in PAO1 (**Figs. 3G, 3H, 4G, and 4H**), indicating that this residue functions as a critical orienting or stabilizing element during cargo loading. Additionally, we identified three closely clustered residues that repeatedly participate in Hcp–effector interfaces (**Fig. 5D**). Although their chemical identities vary across species, they consistently form a polar-enriched binding cleft, pointing to a structurally conserved interaction hotspot that facilitates effector stabilization within the Hcp lumen. Consistently, mutating the corresponding binding region of Tse2 (Q139E) disrupted its interaction with Hcp1 and abolished T6SS-dependent delivery and intoxication (**Extended Data Figs. 10A and 10B**). Moreover, beyond these broadly conserved structural features, finer interaction features exhibit species-specific tailoring. For example, in PAO1 Hcp1, residues S31, S111, and N124—located on the neighboring subunits adjacent to the E118-mediated electrostatic interface—frequently contact flexible regions of its cognate effectors (i.e., Tse1, Tse2, and Tse4) (**Extended Data Fig. 10C**). A comparable distribution of these local interactions was not observed in the heterologous systems (*S.* Typhi and *S. marcescens*). Collectively, our findings highlight a balance between strict structural conservation and local adaptability for Hcp–cargo recognition. The conserved lumen architecture provides a universal platform for cargo accommodation, while variable local interactions fine-tune binding to individual effectors. The polar-biased, variation-tolerant interaction mode allows Hcp proteins to preserve a core secretion mechanism while maintaining the evolutionary flexibility to engage highly diverse cargo repertoires.

## Discussion

Bacteria employ protein secretion systems to compete with neighboring cells and shape microbial community structure. Although the T6SS can deliver a diverse repertoire of toxic effectors, the mechanisms governing effector recruitment—particularly for cargos carried within the Hcp inner tube during transportation—remain poorly understood. Unlike spike-associated effectors, which are often recognized through specialized adaptors and conserved interaction motifs, many Hcp-associated cargo effectors lack identifiable loading domains. This observation raises the fundamental question of how the Hcp tube selectively accommodates structurally and functionally diverse effectors while preserving the architectural constraints required for secretion. Building on these questions, in this study, we combined DMS, structural analysis, and functional competition assays to better understand the principles underlying Hcp-mediated cargo recognition and delivery. Our findings define Hcp1 as a structurally conserved yet interaction-permissive cargo platform (**Fig. 6**). Rather than relying on highly conserved sequence motifs, Hcp1 engages diverse effectors through dispersed lumen-facing interfaces enriched in sequence-variable residues. These residues enable interface diversification for cargo recognition while maintaining the structural constraints required for T6SS inner-tube assembly and efficient effector delivery. Our observation that cargo-interacting residues are concentrated within evolutionarily variable regions further suggests that Hcp–effector interfaces can evolve without disrupting the core architecture of the secretion system.

**Fig. 6:**
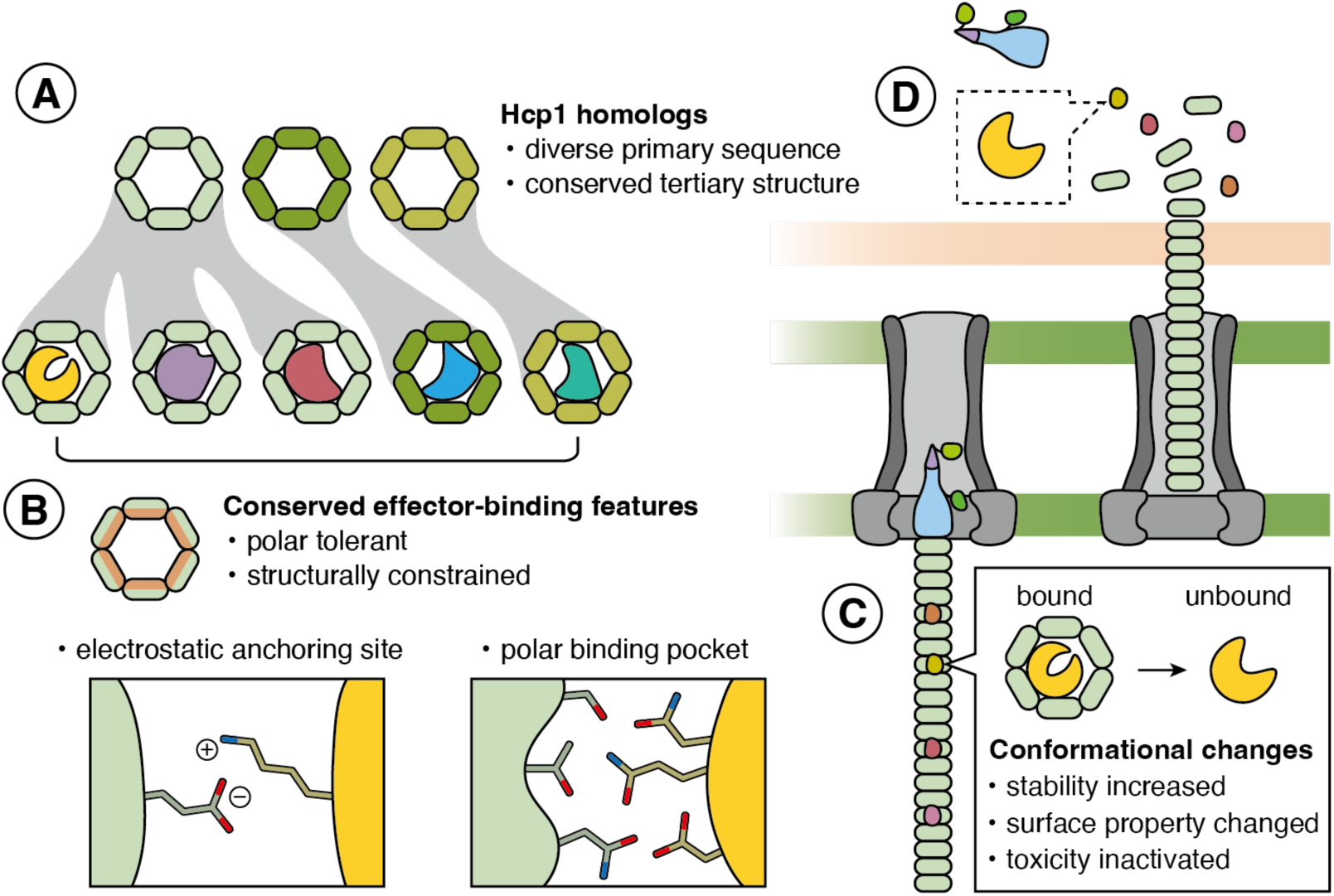
Proposed model of Hcp1-mediated cargo recognition and delivery. (A) Hcp1 functions as a structurally conserved yet interaction-permissive cargo platform. Although cargo-interacting residues diverge across species, the overall architecture of the Hcp1 ring is maintained. Hexameric rings are illustrated with cognate cargo effectors accommodated within the central lumen. (B) Hcp1–cargo recognition relies on conserved lumen-facing structural features, including a recurring electrostatic anchoring site and polar-enriched interaction surfaces. (C) Upon loading into the Hcp1 lumen, effectors may adopt alternative conformations that facilitate stabilization, accommodation, or suppression of premature toxicity. (D) Following T6SS-mediated delivery into the target cell, effectors are released from the Hcp1 tube and regain their active conformations.

A key insight from our work is the nature of Hcp1–cargo interfaces. We observed mostly polar interactions distributed across several contact points, rather than a single high-affinity binding pocket. This interaction surface differs from typical chaperone–substrate systems, which often use hydrophobic motifs and buried surfaces. For example, chaperones belonging to the Hsp70 protein family contain a conserved hydrophobic cleft for client binding [43]. In the case of Hcp1, Hcp1-mediated cargo recognition balances specificity and adaptability through multiple, partially redundant interactions, potentially enabling bacteria to diversify their effector sets while still using a shared secretion system. Furthermore, we consistently observed that the Hcp1 residue E118 engaged in electrostatic contacts across all analyzed Hcp–effector complexes. Notably, electrostatic interactions have been shown to facilitate initial client attraction in the bacterial chaperone Spy [44, 45]. Together, these observations indicate that E118 may function as an early anchoring site during cargo recognition by Hcp1. Future studies will be required to determine if E118 indeed directly contributes to the initial stages of cargo engagement by Hcp1.

Expanding on the adaptable nature of the Hcp1 lumen, our analyses support that Hcp1 may play a more dynamic role in cargo accommodation than previously appreciated. The distinct conformations we observed for Tse2 and Tse4 in their Hcp1-bound states, relative to unbound or predicted structures, suggest that Hcp1 facilitates the stabilization or remodeling of effectors into delivery-competent conformations within the confined lumen environment (**Fig. 6**). This phenomenon may represent a general feature of Hcp-mediated cargo loading. Consistent with this idea, cryo-EM analysis of the *Bacteroides fragilis* T6SS revealed that the effector Bte1 undergoes conformational rearrangements to fit within the Hcp lumen [46]. Moreover, we found that different effectors exhibited distinct dependencies on Hcp1-mediated stabilization in our study, implying that individual cargos impose unique structural and biophysical requirements during the loading process. Together, these observations provide evidence that Hcp functions not merely as a passive conduit but as a dynamic scaffold capable of accommodating chemically and structurally diverse substrates. The biological significance of these conformational changes remains unclear. One possibility is that Hcp-mediated remodeling suppresses effector activity within producer cells prior to secretion, thereby preventing self-intoxication. Although direct visualization of cargo remodeling during loading and secretion will be required to test this hypothesis, our findings establish a framework linking Hcp-mediated cargo accommodation to effector deployment during interbacterial antagonism.

Deep mutational scanning (DMS) has emerged as a powerful approach for dissecting proteins involved in complex biological functions and interactions. By applying DMS to Hcp1 under physiologically relevant interbacterial competition conditions, we generated a comprehensive residue-level functional landscape of Hcp1 activity. Unsupervised clustering of mutational response profiles further partitioned Hcp1 residues into distinct functional groups characterized by different substitution preferences. Based on these results, we identified two clusters strongly associated with effector interactions. One exhibited high tolerance to polar substitutions, underscoring the importance of polar contacts in cargo recognition, whereas the other was broadly mutation-tolerant but particularly sensitive to proline substitutions, suggesting that structural integrity is critical for accommodating effectors within the Hcp lumen. Interestingly, these interaction-associated features appear to extend beyond *P. aeruginosa*. In both *S.* Typhi and *S. marcescens*, Hcp residues involved in effector interactions mapped to positions corresponding to the same two functional clusters identified in Hcp1, indicating that common physicochemical principles may govern Hcp-mediated cargo recognition across diverse bacterial species despite substantial sequence divergence. Beyond providing new insights into Hcp function, our study also highlights the utility of competition-coupled DMS as a general strategy for investigating bacterial secretion systems. By linking residue-level protein variation directly to bacterial competitive fitness, this approach enables functional dissection of secretion machineries under biologically relevant conditions. As many of the activities of secretion systems are only fully manifested during cell–cell interactions, competition-based selection offers a powerful framework for resolving the functional architecture of complex bacterial nanomachines. More broadly, this strategy should be applicable to a wide range of secretion systems and contact-dependent antagonistic pathways.

Tse3 proved to be a notable exception in our analysis. Although we successfully characterized Hcp1’s interactions with Tse1, Tse2, and Tse4 using either experimentally determined or high-confidence structural models, we were unable to obtain sufficient structural information for the Hcp1–Tse3 complex. Attempts to purify the complex were unsuccessful, and AlphaFold3 predictions yielded consistently low confidence scores, precluding reliable analysis of the interaction interface. Interestingly, Tse3 is substantially larger than the other Hcp1-associated effectors examined in this study, with a molecular weight of approximately 44.4 kDa. Thus, it is possible that Tse3 may not be accommodated in Hcp1 ring through the same loading mode used by smaller cargo effectors. Instead, Tse3 may undergo more extensive conformational remodeling than other Hcp cargoes during loading. Alternatively, cargo accommodation may involve interactions extending beyond a single Hcp ring, thereby increasing the available loading volume. Although these possibilities require further validation, they highlight the potential diversity of cargo-loading strategies employed by Hcp proteins. Future structural studies of the Hcp1–Tse3 complex will be required to determine if large effectors utilize distinct mechanisms for cargo accommodation and delivery.

The dynamics of Hcp-mediated cargo loading remain poorly understood. Although our structures provide snapshots of cargo-bound Hcp1 complexes, they offer limited insight into the process by which effectors are recruited, loaded, and accommodated within the Hcp lumen. Interestingly, a recent study on the *P. aeruginosa* H3-T6SS proposed a sequential cargo-loading mechanism, whereby Hcp3 progressively assembles around its cognate effector Tce1 [47]. Whether this model is broadly conserved among Hcp proteins remains unclear, particularly given that Hcp1 and Hcp3 belong to distinct evolutionary subgroups and are associated with secretion systems that deploy different effector repertoires and biological functions [48–50]. We did not identify particle classes corresponding to partially assembled loading intermediates from our cryo-EM analyses of the Hcp1–Tse2 and Hcp1–Tse4 complexes. However, the absence of such classes does not exclude a sequential assembly pathway, as transient intermediates may be difficult to capture during complex purification or cryo-EM sample preparation. Consequently, future biochemical and structural studies will therefore be required to determine whether Hcp-mediated cargo loading follows a conserved assembly pathway or if distinct Hcp proteins have evolved specialized cargo-loading strategies.

Finally, based on our findings presented herein, we provide a framework for effector discovery and synthetic engineering. The absence of conserved loading motifs has historically hindered the identification of Hcp-associated cargo effectors using sequence-based approaches alone. However, we show that Hcp-mediated cargo recognition instead likely relies on disperse physicochemical compatibility and conserved lumenal geometry. Accordingly, the functional and structural features identified in our study—including permissive interaction surfaces, polar-enriched interfaces, and characteristic mutational tolerance patterns—provide a framework for future identification of candidate Hcp cargos across diverse bacterial species. More broadly, the adaptable nature of the Hcp lumen raises the possibility of engineering programmable T6SS-based delivery systems. For example, our observation that Tse2 engages Hcp1 through discrete structural elements indicates that minimal loading determinants could potentially be fused to heterologous proteins for synthetic cargo delivery. Future studies integrating functional screening, structural analysis, and engineered cargo systems will help clarify how substrate loading, translocation, and release are coordinated during T6SS-mediated transport.

## Methods

### Bacteria and culture conditions

The strains used in this study are detailed in **Supplementary Table 3**. We used the following *Escherichia coli* strains: DH5α in cloning and plasmid maintenance; S17-1 λ pir for conjugal transfer of pEXG2 plasmid into *Pseudomonas aeruginosa*; BL21 (DE3) pRIL and Rosetta 2 for overproduction and protein purification. *P. aeruginosa* PAO1 was used for library construction, protein secretion assays, and interbacterial competition assays. *Enterobacter cloacae* ATCC 13047 was used for interbacterial competition assays. All bacteria were routinely cultured at 37 °C in Lysogeny Broth (LB) medium, unless otherwise noted. Antibiotics and chemicals were used at the following concentrations: 50 μg/mL streptomycin; 150 μg/mL carbenicillin; 50 μg/mL kanamycin; 25 μg/mL chloramphenicol; 15 μg/mL gentamicin; 100 μg/mL trimethoprim; 25 μg/mL irgason; 1% (w/v) L-(+)-rhamnose; and 200 mM sucrose.

### Plasmid construction

All plasmids and primers used in this study are listed in **Supplementary Tables 4 and 5**. Chromosomal mutations in *P. aeruginosa* PAO1 were generated using the pEXG2 vector [51]. For gene deletions, 800-basepair (bp) regions flanking the mutation site were amplified by polymerase chain reaction (PCR), and inserted into the vector using Gibson assembly at the XbaI and SacI restriction sites [52]. For site-directed mutagenesis, primers containing the desired mutations were paired with a complementary primer in the opposite orientation to amplify a mutated pEXG2 plasmid copy. To generate a strain expressing an N-terminally VSV-G-tagged Tse4 from its native chromosomal locus, the VSV-G coding sequence was incorporated into the PCR primers used to amplify the *tse4* allele prior to assembly into pEXG2. To insert ampicillin resistance gene at the Tn7 sites of *P. aeruginosa*, the gene was amplified from the pPSV18 plasmid and then introduced into the pEXG2 plasmid via Gibson assembly.

For heterologous expression of Hcp1 in *P. aeruginosa* strains, *hcp1* (PA0085) was amplified and cloned into the SacI and XbaI restriction sites of the pPSV39 vector. To purify the Hcp1-Tse2^NT^(T79A/S80A) and Hcp1-Tse4 complexes, the corresponding genes were cloned into separate plasmids. First, *hcp1* was amplified and cloned into the NdeI and XhoI restriction sites of the pET29b(+) vector to include a C-terminal Strep-Tactin tag (WSHPQFEK), yielding pET29b(+)_Hcp1-strep. Next, *tse2* (PA2072; T79A/S80A) and *tse4* (PA2774) were cloned into the multiple cloning site (MCS)-1 of the pCDFDuet-1 vector at NcoI/NotI and BamHI/NotI sites, respectively, resulting in the pCDFDuet-1_Tse2_His and pCDFDuet-1_Tse4_His plasmids for the subsequent experiments. For Tse2 stability assay, primers were designed to amplify *tse2* (T79A/S80A) fused with a N-terminal hexa-histidine tag, then the amplicon was cloned into pSCrhaB2 vector. Single-point mutations in *hcp1* were introduced into pET29b(+)_Hcp1-Strep via inverse PCR using primers carrying the corresponding mutation.

### Genetic manipulation

For *P. aeruginosa* strain generation, pEXG2 mutation constructs were transformed into *E. coli* S17-1 λpir as donor strains. The donors were then mixed and incubated with *P. aeruginosa* recipients on LB agar plates at a 1000:1 donor-to-recipient ratio. The cell mixtures were incubated for 6 hours at 37°C to facilitate conjugation. Afterwards, the mixtures were scraped, resuspended in 1 mL LB medium, and plated on LB agar plates containing irgasan and gentamicin to select for cells with the mutant construct inserted into the chromosome. The following day, single colonies were inoculated into 3 mL of nonselective LB medium, shaken at 37°C for 6 hours, then counterselected on LB no-salt (LBNS) agar plates supplemented with sucrose. Gentamicin-sensitive, sucrose-resistant colonies were screened for allelic replacement by colony PCR, and mutations were confirmed through Sanger sequencing of the PCR products.

### Bacterial competition assays

For inter-strain competition assays, *E. cloacae* served as the attacker and *P. aeruginosa* strains as the recipients. For intra-strain competition assays, *P. aeruginosa* Δ*retS* strains were used as attackers, whereas *P. aeruginosa* strains lacking specific effector–immunity pairs (*tse1*/*tsi1*, *tse2*/*tsi2*, and/or *tse4*/*tsi4*) served as recipients. Strains were grown overnight, and then cells were harvested by centrifugation. Attacker and recipient strains were mixed at an appropriate donor-to-recipient ratio. Competition assays were initiated by spotting three 5 μL aliquots of each mixture onto nitrocellulose filters placed on LBNS agar plates containing 3% (w/v) agar. Plates were incubated at 37 °C for 6 hours. After incubation, cells were recovered by scraping individual spots from excised nitrocellulose filter sections into LB medium. The suspensions were serially diluted and plated on selective media to enumerate colony-forming units (CFUs). The competitive index was calculated as the ratio of the attacker-to-recipient output ratio to the attacker-to-recipient input ratio.

### Library construction

To construct the Hcp1 library for deep mutational scanning, 86 sites on *hcp1* were mutagenized, with each site mutated to all desired amino acids. GenScript synthesized DNA oligonucleotide libraries on a programmable column. The pooled mutagenesis library was subsequently cloned into the pPSV39 backbone (SacI/XbaI) and transformed into *P. aeruginosa* PAO1 Δ*hcp1* through electroporation, resulting in a collection of >170,000 colonies stored in LB broth with 10% glycerol at −80 °C.

### Functional screening

The screen was conducted using the interbacterial competition assay described above, with minor modifications. *E. cloacae* served as the attacker strain, while the PAO1 Hcp1 library acted as the recipient. The overnight-grown attacker was diluted to optical density (OD)_600_ = 0.05 in 25 mL LB medium. For the recipient, the stored library was thawed and adjusted to OD_600_ = 0.05 in 25 mL LB medium supplemented with gentamycin and 0.5 mM IPTG. The cultures were then shaken at 37 °C for 4 hours for induction. Then, cells were collected and washed once with LB. Attacker and recipient strains were mixed at a 3:1 OD ratio, and then five 5 μL aliquots of each mixture were spotted onto nitrocellulose filters placed on LBNS agar plates containing 3% (*w/v*) agar and 0.5 mM IPTG. After 7 hours of incubation, the cells were recovered by scraping individual spots from excised nitrocellulose filter sections. The cells were resuspended and diluted with LB medium, and then plated on selective media containing irgasan and gentamicin. All grown colonies (>450,000 clones) were harvested and stored for high-throughput sequencing.

### Determination of the functionality of Hcp1 variants within the library

To assess the functional impact of hcp1 amino acid substitutions during interbacterial competition, we harvested cells and extracted plasmid DNA for targeted amplicon sequencing as described previously [53]. In brief, 10 ng of the extracted plasmid DNA was used as the template for a first-round PCR amplification (18 cycles) of the *hcp1* gene region using target-specific primers with adapter overhangs (listed in **Supplementary Table 5**). Amplicons of the expected size (∼561 bp) were gel-purified using a QIAquick Gel Extraction Kit. To generate sequencing libraries, 10 ng of the purified first-round PCR products was subjected to a second-round PCR (8 cycles) using KAPA HiFi HotStart ReadyMix to incorporate Illumina i5 and i7 index sequences (Illumina DNA/RNA UD Indexes Set D, Tagmentation). The indexed amplicons were subsequently purified using SPRI beads (Beckman Coulter), quantified, and validated for the expected library fragment size with Qubit and Agilent 5200 Fragment Analyzer. The individual libraries were then normalized to 2 nM, pooled, and quantified using the KAPA Library Quantification Kit for Illumina Sequencing Platforms. Sequencing was performed using 300-bp paired-end reads on Illumina MiSeq i100 (15% PhiX spike-in).

Raw paired-end sequencing reads provided in FASTQ format were processed using a custom C++ analysis pipeline utilizing the SeqAn sequence analysis library. To isolate the targeted mutation region, the script scanned each read for the upstream plasmid flanking sequence pattern “ATCTGTATATCTAGA”. Upon detection, the immediate downstream 450-bp sequence region was extracted. Since the raw reads were sequenced in the reverse-complement orientation relative to the *hcp1* coding sequence, the extracted fragments were converted to their true 5’-to-3’ reverse-complement sequences to restore the original gene coding frame. The corrected DNA segments were subsequently translated *in silico* into primary protein sequences using the standard genetic code.

To minimize the impact of sequencing artifacts, unique DNA and protein variants were compiled into frequency tables, and only high-confidence variants with an absolute read abundance threshold of ≥ 5 counts were preserved for downstream analysis. High-confidence variant sequences were globally aligned against wild-type *hcp1* nucleotide and amino acid reference templates. Mutation detection was conducted to systematically log substitutions, insertions, deletions (indels), or premature stop codons.

Functional scores for each Hcp1 variant (mut) were calculated as the relative survival after competition, normalized to wild-type Hcp1 (wt), using the following equation:

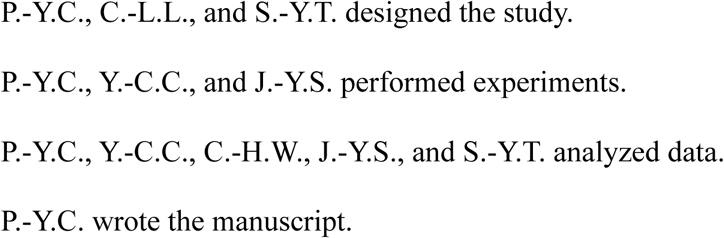

For downstream analysis, the overall score distribution of the DMS library was visualized via kernel density estimation (KDE) plots generated using Python (v3.11) with the Seaborn and Matplotlib packages.

### Unsupervised learning

To identify functional clusters of Hcp1 residues, we performed unsupervised learning using Uniform Manifold Approximation and Projection (UMAP) following a general framework described previously [38]. First, amino acid substitutions were grouped into eight physicochemical classes: positively charged [(+) ; R, H, K], negatively charged [(-) ; D, E], aromatic (F, W, Y), amide (N, Q), nucleophilic (C, S, T), hydrophobic (I, L, V, M), small (G, A), and proline (P). For each residue position, functional scores were averaged within each amino acid class. A design matrix was then constructed in which rows represented Hcp1 residue positions and columns represented the average functional score of each amino acid substitution, calculated as the mean across three biological replicates. UMAP was then applied to this matrix using the cosine distance metric with the following hyperparameters: n_neighbors = 8, n_components = 2, min_dist = 0, and n_epochs = 500 (uwot R package, random seed = 23). To identify discrete functional modules, HDBSCAN was subsequently applied to the two-dimensional UMAP embedding (minPts = 4; dbscan R package), assigning each position to one of six clusters. All analyses were performed in R v4.5.0.

### Protein purification

For protein expression, *E. coli* BL21(DE3) Rosetta 2 cells were co-transformed with the Hcp1-expressing plasmid (pET29b(+)_Hcp1-strep) and one of the toxin-expressing plasmids (pCDFDuet-1_Tse2_His or pCDFDuet-1_Tse4_His). Cells were selected on LB agar supplemented with chloramphenicol, kanamycin, and streptomycin. Cultures were grown in 2xYT medium at 37 °C with 200 rpm shaking until they reached mid-log phase, at which point expression was induced with 1 mM IPTG for 4 hours. Cells were harvested by centrifugation (4,000 × *g*, 20 minutes, 4 °C) using a JLA-8.1000 rotor (Beckman Coulter), flash-frozen in liquid nitrogen, and stored at -80 °C. Cell pellets were resuspended in lysis buffer (20 mM Tris-HCl, pH 7.5; 300 mM NaCl; 10 mM imidazole, pH 7.0; 1 mM DTT; 1 mg/mL lysozyme; and a protease inhibitor cocktail). Cells were lysed by sonication (10 cycles of 10 s at 50% amplitude with 30 s cooling intervals). The lysate was cleared by centrifugation (20,000 × *g*, 30 minutes, 4 °C) using a JA-25.50 rotor (Beckman Coulter). The cleared lysate was first purified by gravity-flow Ni-NTA affinity chromatography (2 mL resin volume, Qiagen). Bound proteins were eluted with a linear imidazole gradient up to 300 mM. Eluted fractions containing the complex were pooled and further purified using a 1 mL Strep-Tactin Sepharose column (IBA). The Hcp1-Tse2 complex was eluted with 4.5 mL of Tris-elution buffer (100 mM Tris-HCl, pH 7.5; 150 mM NaCl; 2.5 mM d-desthiobiotin). In contrast, the Hcp1-Tse4 complex was eluted with 4.5 mL of Hepes-elution buffer (20 mM Na-Hepes, pH 7.4; 150 mM NaCl; 2.5 mM d-desthiobiotin) for a subsequent crosslinking reaction. Protein purity was verified by 15% SDS-PAGE with Coomassie Brilliant Blue staining.

The purified Hcp1-Tse4 protein complex was diluted to 1 mg/mL and crosslinked with a final concentration of 0.125% glutaraldehyde (Sigma-Aldrich). The reaction was performed on ice for 5 minutes and quenched with 1 M Tris-HCl, pH 7.5. The crosslinked complex was then subjected to size exclusion chromatography (SEC) using a Superdex 200 10/300 Increase GL column (Cytiva) equilibrated with SEC buffer (20 mM Tris-HCl, pH 7.5; 300 mM NaCl). Eluted fractions were analyzed by 15% SDS-PAGE, and peak fractions were pooled and concentrated using a 30 kDa MWCO Amicon concentrator.

### Cryo-EM sample preparation, data collection and data processing

For grid preparation, proteins were diluted to 0.15 mg/mL in a sample buffer (20 mM Tris-HCl, 300 mM NaCl, 0.01% Tween-20) adjusted to either pH 5.0 for the Hcp1-Tse2 complex or pH 8.0 for the Hcp1-Tse4 complex. A 4 µL aliquot of the sample was applied to glow-discharged Quantifoil R2/1 200-mesh copper grids (Quantifoil Micro Tools GmbH) coated with a 2 nm continuous carbon film. The grids were then blotted for 4 s (blot force 0) and plunge-frozen into liquid ethane using a Vitrobot Mark IV system (Thermo Fisher Scientific) set to 4 °C and 100% humidity. Grids were first screened on a 200 keV Talos Arctica microscope (Thermo Fisher Scientific) equipped with a Falcon III detector operating in linear mode. Screening images were recorded at 120,000× magnification (pixel size 0.86 Å/pixel) with a defocus of −2.5 µm. High-resolution data were collected on a 300 keV Titan Krios microscope (Thermo Fisher Scientific) equipped with a K3 detector and a GIF Bio-Quantum energy filter (Gatan). Data were acquired in super-resolution mode using EPU-3.6.0 software at a magnification of 105,000×, corresponding to a physical pixel size of 0.83 Å/pixel (0.415 Å/pixel in super-resolution). Movies were recorded with a defocus range of −1.0 to −1.6 µm. The energy filter slit width was set to 15 eV. Each movie consisted of 50 frames with an exposure time of 1.34 s, resulting in a total dose of ∼49.8 e−/Å² (∼0.996 e−/Å² per frame). Acquired super-resolution movies were motion-corrected, dose-weighted, and binned twofold using MotionCor2 (7 × 5 patch), resulting in a final pixel size of 0.83 Å/pixel. Motion-corrected micrographs were imported into cryoSPARC v4.5.1 for subsequent processing. CTF estimation was performed using the CTF estimation job (CTFFIND4) [54].

Particles were extracted with a 256-pixel box size and subjected to multiple rounds of 2D classification. Particles from selected 2D classes were used to generate ab initio models. Iterative 3D heterogeneous refinement was then performed to identify distinct conformational states and to separate toxin-bound (Hcp1-Tse2, Hcp1-Tse4) from unbound (Hcp1-apo) complexes, which likely resulted from spontaneous toxin dissociation during purification. The remaining particles were further refined using Non-Uniform Refinement (NU-refinement) without symmetry constraints. The initial reconstructions and micrographs revealed that double layers exhibited significant structural flexibility, thereby preventing high-resolution alignment of the full complex. This outcome is consistent with the observation that a stable double-layer structure cannot be resolved for the isolated Hcp1 ring either. To improve the local resolution and clarity of specific regions, focused refinement was applied using a soft mask.

The final composite map was generated by combining the maps from focused refinement. Final maps were sharpened, and resolutions were estimated within cryoSPARC [55]. The overall resolution was estimated using the Fourier Shell Correlation (FSC) 0.143 criterion, and then local resolution was calculated within cryoSPARC. Density maps were visualized with UCSF ChimeraX [56]. Detailed information on the reconstruction process is provided in **Extended Data Figs. 4 and 7**, with statistical parameters summarized in **Supplementary Table 6**. Atomic models of the Hcp1-Tse2, Hcp1-Tse4, and Hcp1-apo complexes were built using an AlphaFold3-predicted model as a starting template. Model refinement was performed iteratively using Coot [57] and PHENIX[58]. Summary statistics for the final models are presented in **Supplementary Table 6**.

To characterize the interbacterial interaction interfaces within the protein complex, atomic contacts between the Hcp1 subunits and effectors were analyzed using UCSF ChimeraX. The resulting interaction network was visualized as a chord diagram using the circlize package in R.

### Molecular dynamics simulation of the hcp1 tubular complex

A triple-layer of the Hcp1 tubular model was generated using the Vibrio cholerae sheath/tube complex (PDB ID 5OJQ) as a structural scaffold. The sheath components were removed from the template, and the hexameric Hcp1 rings determined in this study were aligned onto the remaining tube coordinates to establish the vertical stacking arrangement. Relative orientation and packing of the rings were further refined by optimizing the electrostatic complementarity between the anionic and cationic faces of adjacent subunits. The resulting tubular complex was parameterized using the AMBER ff14SB [59] force field and solvated in a periodic box of TIP3P water molecules with a minimum buffer of 8.0 Å. The system was neutralized by adding 18 Na+ counterions. Energy minimization was performed in three stages, using both the steepest-descent and conjugate-gradient algorithms to remove steric clashes. Following minimization, the solvated complex was heated to 300 K and equilibrated in the isothermal-isobaric (NPT) ensemble. A Langevin thermostat was used to maintain the temperature at a collision frequency of 1 ps^-1^. All covalent bonds involving hydrogen atoms were constrained using the SHAKE algorithm, and a conservative integration time step of 1 fs was employed. Production simulations were conducted under NPT conditions at 300 K and 1.0 atm for a total duration of 50 ns.

Atomic coordinates were recorded every 1 ps for subsequent structural analysis and the calculation of time-averaged properties.

### Tse2 stability assays

Stability assays were conducted as previously described [24] with slight modifications. In brief, pET29b plasmids encoding individual Hcp1 variants and pSCRhaB2 carrying *tse2*^NT^ were co-transformed into *E. coli* BL21 (DE3) pRIL. Overnight cultures were diluted into 25 mL LB medium to an OD_600_ of 0.05 and grown to log phase. Protein expression was induced with 0.5% rhamnose and 0.5 mM IPTG, and cultures were harvested after 4 hours. For each sample, cells equivalent to 1 OD unit were collected, resuspended in 2× SDS sample buffer, boiled, and analyzed by immunoblotting.

### Protein secretion assays

*P. aeruginosa* strains were cultured overnight and diluted into 25 mL LB medium to an OD_600_ of 0.05. Cultures were grown to logarithmic phase, then cell pellets and culture supernatants were collected. For intracellular protein samples, cell pellets were resuspended in 2× SDS sample buffer. To collect secreted proteins, 1.5 mL of culture supernatant was first filtered through a 0.22 μm filter. Deoxycholic acid was then added to a final volume of 45 μL (1%), and the mixture was incubated at room temperature for 10 minutes. Subsequently, 225 μL of 100% (w/v) trichloroacetic acid was added to precipitate proteins. Samples were incubated overnight at −20 °C, thawed, and centrifuged to pellet the precipitated proteins. The supernatant was removed, and the pellet was resuspended in 100 μL of 1 M Tris supplemented with 1× SDS sample buffer. Samples were sonicated for 20 minutes to fully resuspend the proteins. Finally, intracellular and secreted protein samples were analyzed by immunoblotting.

### Immunoblotting assays

To analyze the levels of intracellular and secreted proteins, samples were boiled at for 10 minutes and then equal volumes were loaded for SDS-PAGE. After being transferred to nitrocellulose membranes, membranes were blocked in TBST (10 mM Tris-HCl pH 7.5; 150 mM NaCl; 0.1% (w/v) Tween-20) with 3% (w/v) BSA for 30 min at room temperature, before being incubated for 1 hour at room temperature with primary antibodies (anti-Hcp1, anti-Tse1, anti-Tse2, anti-VSV-G, or anti-RpoB) diluted to an appropriate concentration in TBST with 0.1% BSA. Blots were then washed by TBST, followed by incubation for 30 minutes at room temperature with the secondary antibodies (anti-mouse HRP-conjugated IgG or anti-rabbit HRP-conjugated IgG) diluted 1:5000 in TBST with 0.1% BSA. Finally, blots were washed again by TBST, developed, and visualized using a UVP Biospectrum 815 system.

### Hcp protein sequence conservation analysis

To assess Hcp1 sequence conservation at the species level, 1,809 homologous sequences were retrieved from the NCBI RefSeq representative prokaryotic genomes database using the *P. aeruginosa* PAO1 Hcp1 protein sequence as a query in tBLASTn searches. For strain-level analysis, Hcp1 homologs within *P. aeruginosa* were identified using NCBI BLASTp. Retrieved 1,633 sequences were aligned using ClustalW, and sequence conservation was quantified using Jensen–Shannon divergence [60]. The sequence logos of Hcp1 were generated with WebLogo [61].

## Supporting information

Supplementary Table 1: P. aeruginosa PAO1 Hcp1 residues in 6 UMAP-defined clusters

Supplementary Table 2: S. Typhi Hcp and S. marcesens residues in 6 UMAP-defined clusters

Supplementary table 3: Bacterial strains used in this study

Supplementary table 4: Plasmids used in this study

Supplementary table 5: Oligonucleotides used in this study

Supplementary Table 6. Cryo-EM data collection, refinement and validation statistics

## Data Availability

The coordinates and structure factors for the Hcp1-apo, Hcp1-Tse2, and Hcp1-Tse4 complexes have been deposited in the Protein Data Bank, with accession codes 24AK, 24AL, and 24AM, respectively. Sequence data associated with this study is available from the Sequence Read Archive at GSE334501.

## Acknowledgements

We thank Dr. Meng-Chiao Ho and the Ting laboratory members for helpful discussions and critical reading of the manuscript; Drs. Joseph Mougous, Kuo-Chiang Hsia, and Hung-Ta Chen for sharing reagents; Drs. Shu-Yun Tung, Hsin-Nan Lin, Chen-Hsin Yu, Po-Pang Chen and Li-Kang Sung for technical assistance; and Dr. John O’Brien for manuscript editing. The cryo-EM experiments were performed at the Academia Sinica Cryo-EM Facility (ASCEM), and the cryo-EM data were processed at the Academia Sinica Grid-computing Center (ASGC). We acknowledge funding support from Academia Sinica (AS-CDA-112-L05 and AS-GCS-115-L07 to S.-Y.T., AS-PD-1151-M09-2 to J.-Y.S., AS-CFII-114A-11 to ASGC, AS-CFII-111-210 to ASCEM) and the National Science and Technology Council, Taiwan (113-2628-B-001-008-MY3 to S.-Y.T.).

## Author Contributions

P.-Y.C., C.-L.L., and S.-Y.T. designed the study.

P.-Y.C., Y.-C.C., and J.-Y.S. performed experiments.

P.-Y.C., Y.-C.C., C.-H.W., J.-Y.S., and S.-Y.T. analyzed data.

P.-Y.C. wrote the manuscript.

## Extended Data

**Extended Data Fig. 1:**
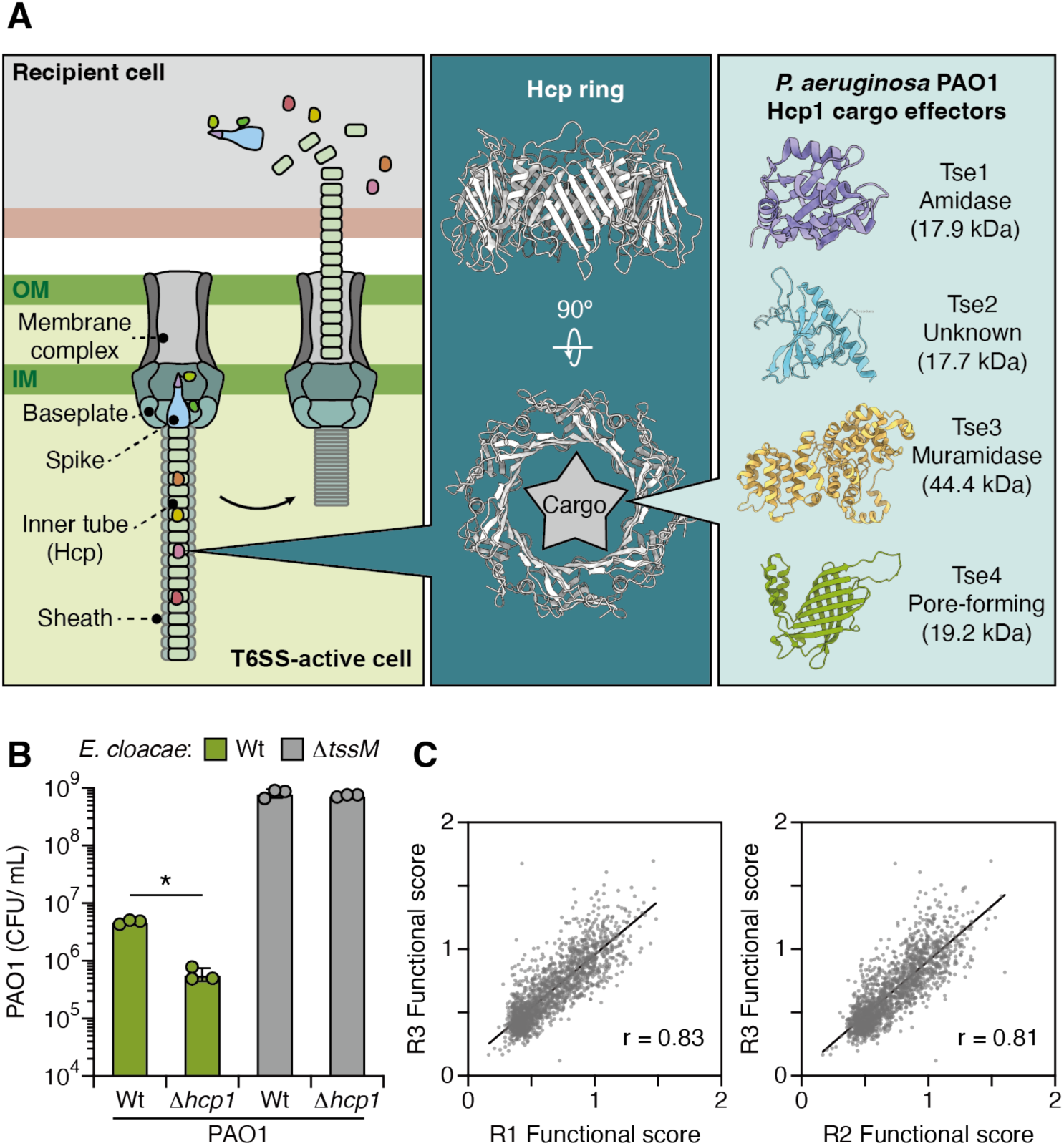
Design of Hcp1 functional screen. (A) Schematic of T6SS-mediated attack during interbacterial competition (left). The inner-tube of T6SS is composed by stacked hexameric Hcp1 ring (middle). In PAO1 H1-T6SS, four effectors are reported to loaded within Hcp1 rings. Structure of Tse1 (PDB: 4FGE), Tse2 (PBD: 5AKO), Tse3 (PDB: 4M5E), and Tse4 (AlphaFold3 model, pTM = 0.71) (B) Survival of PAO1 strains after 6-hour co-incubated with wild-type or Δ*tssM E. cloacae*. Data are presented as means ± SD. n = 3 biological replicates with three technical replicates each. Asterisks indicate statistically significant differences between the indicated mean values (p < 0.05, two-tailed t-test). (C) Correlation between three biological replicates. Each point represents a single variant. Data correlated to Figure 1F.

**Extended Data Fig. 2:**
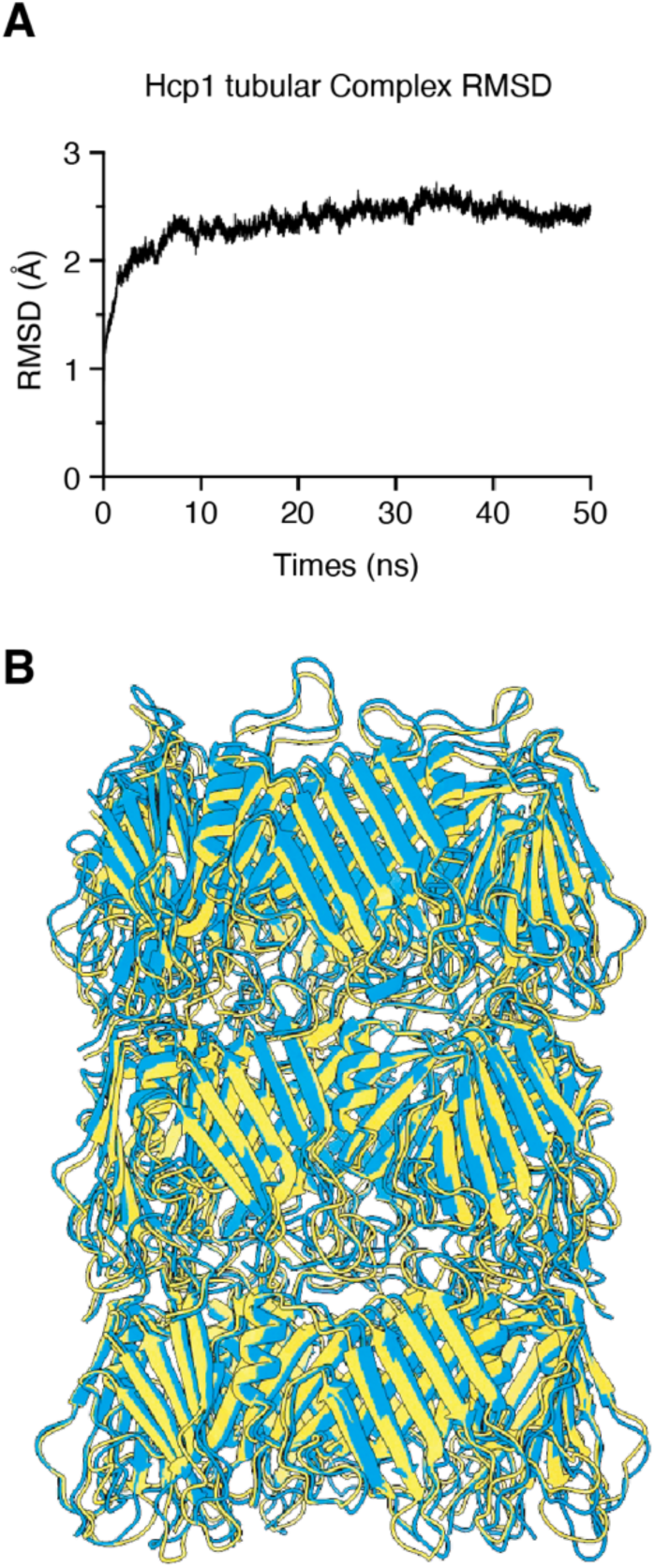
Structural stability and relaxation of the Hcp1 tubular model. (A) Root mean square deviation (RMSD) of the backbone atoms over the 50 ns simulation, showing structural convergence. (B) Superposition of the Hcp1 triple-layer model before (yellow) and after (cyan) MD relaxation. The high degree of structural overlap indicates that the initial docked assembly represents a stable conformation.

**Extended Data Fig. 3:**
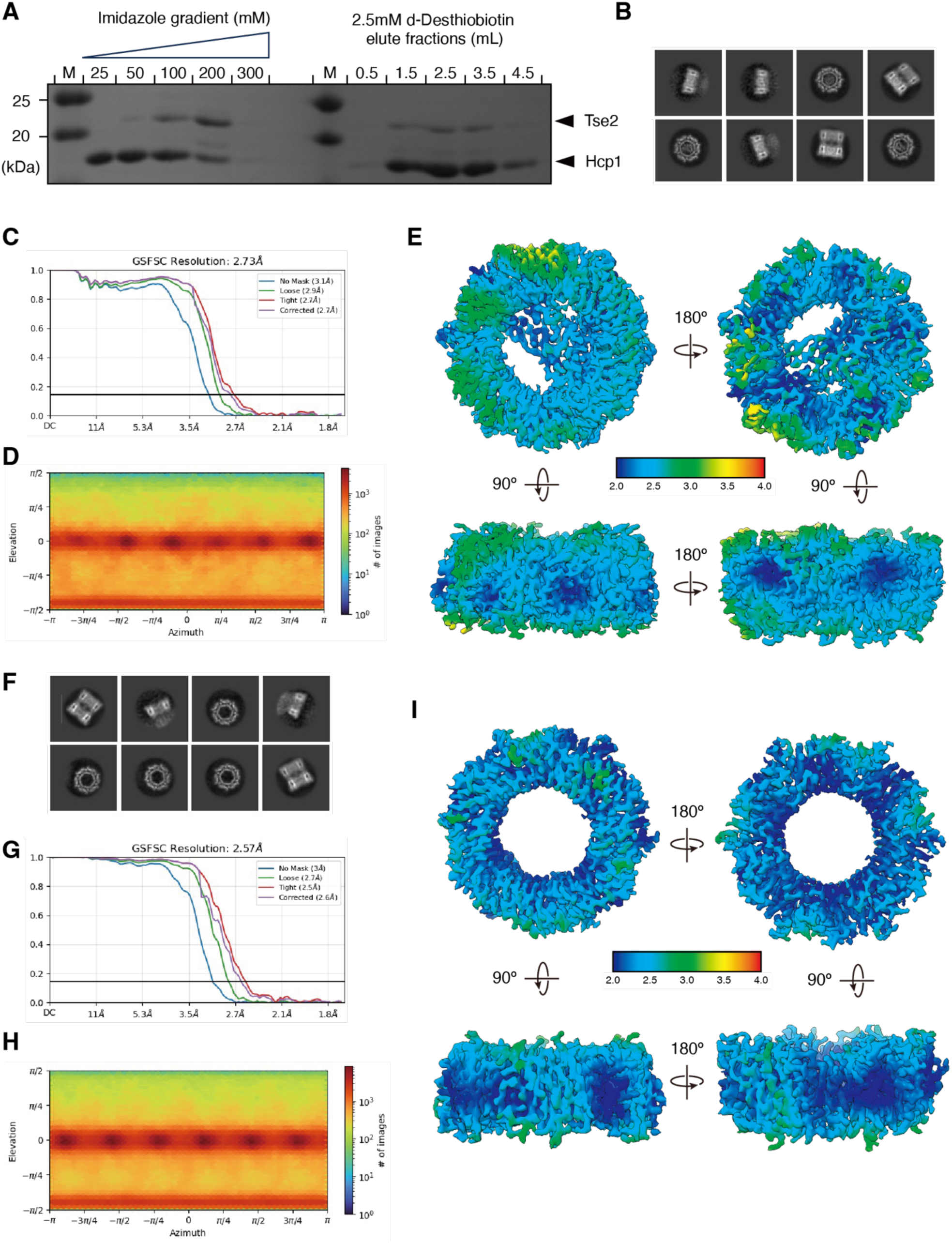
Biochemical purification, validation, and Cryo-EM characterization of the Hcp1-Tse2 and Hcp1-apo complexes. (A) SDS-PAGE analysis of Hcp1-Tse2 purification. Left panels show elution from Ni-NTA resin via an imidazole gradient (mM); right panels show elution from Strep-Tactin resin using d-Desthiobiotin (mL). Molecular weight markers (M) are indicated in kDa. (B-E) Cryo-EM characterization of the Hcp1-Tse2 complex. (B) Representative 2D class averages of Hcp1-Tse2. (C) Gold-standard FSC curve indicating a global resolution of 2.73 Å. (D) Euler angle distribution map for the Hcp1-Tse2 dataset. (E) Cryo-EM density map viewed from the top and side, colored by local resolution (Å). (F-I) Cryo-EM characterization of the Hcp1-apo complex. (F) Representative 2D class averages of the Hcp1-Apo complex. (G) Gold-standard FSC curve indicating a global resolution of 2.57 Å. (H) Euler angle distribution of the particles included in the final 3D reconstruction. (I) Cryo-EM density map viewed from the top and side, colored by local resolution (Å)

**Extended Data Fig. 4:**
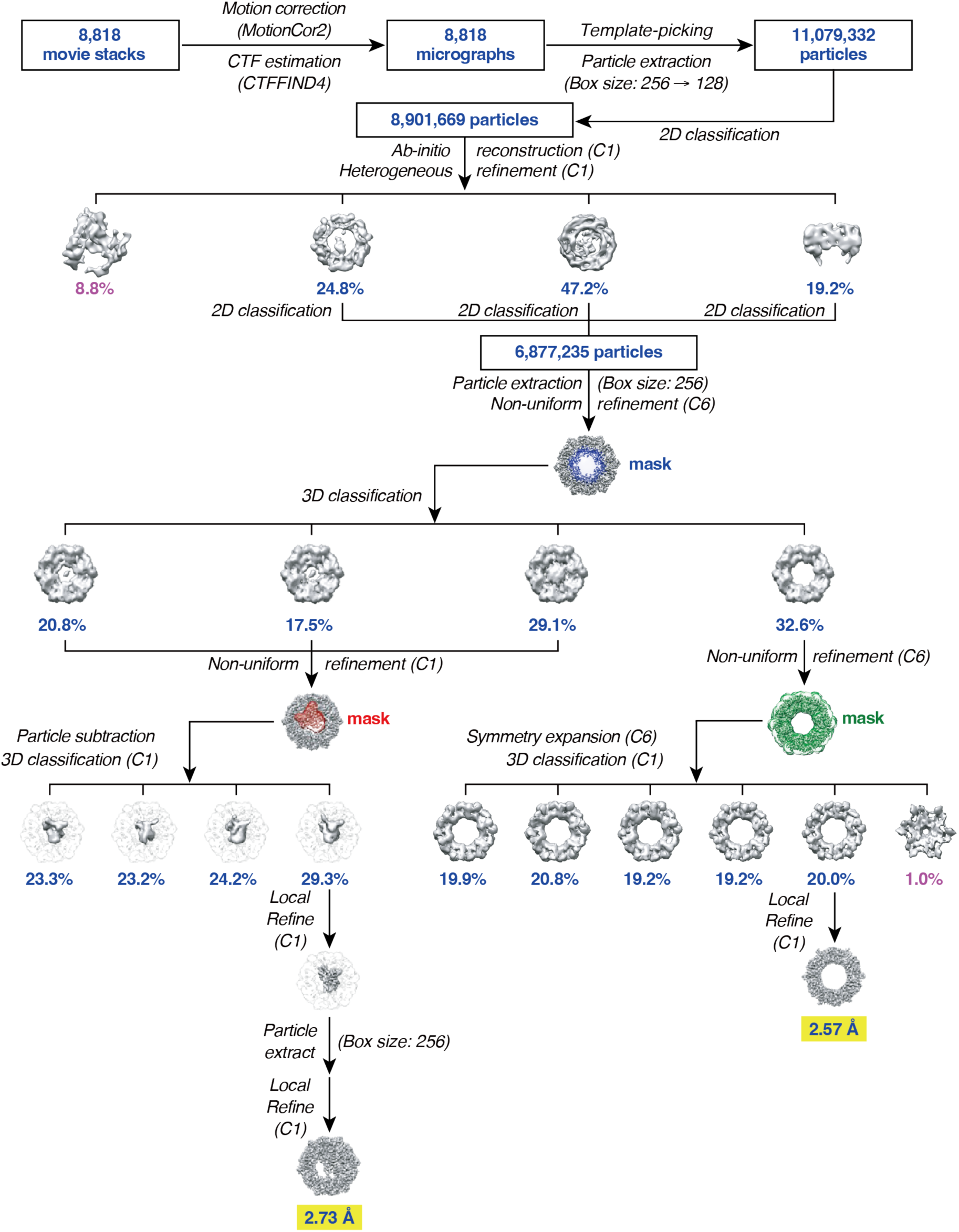
Cryo-EM data processing workflow for the Hcp1-Tse2 complex. Schematic overview of the image processing pipeline for the Hcp1-Tse2 complex. Raw movie stacks (8,818) were pre-processed via patch motion correction and CTF estimation. Particles (11,079,332) were initially template-picked, extracted, and subjected to standard iterative 2D classification. Heterogeneous refinement was then employed, leading to a major 3D class containing 6,877,235 particles. The workflow then split into two concurrent strategies to optimize both global and local resolution: 1) Global Refinement: C6 symmetry was applied during non-uniform refinement. This was followed by final masked (green) local refinement, resulting in a 2.57 Å reconstruction of the hexameric ring. 2) Tse2-Specific Refinement: Particle subtraction (masked in red) was used during focused 3D classification, followed by local refinement. A final particle extraction (256-pixel box) and masked local refinement yielded a 2.73 Å reconstruction, optimizing the visual clarity of the Tse2 component.

**Extended Data Fig. 5:**
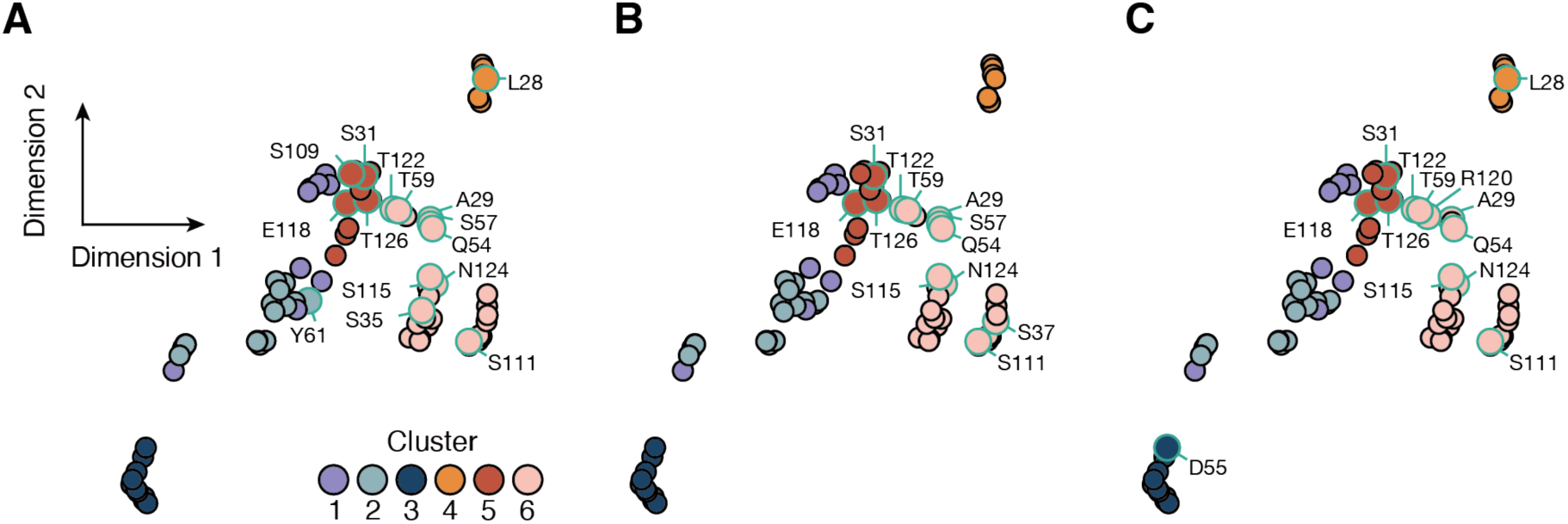
Hcp1 residues interacting with effectors are enriched within two UMAP-defined clusters. Hcp1-Tse2 (A), Hcp1-Tse4 (B), or Hcp1-Tse1 (C) interface residues are mapped to the UMAP projection corresponding to Figure 2A. Each cluster is highlighted with distinct color. Hcp1 residues interacting with effectors are indicated.

**Extended Data Fig. 6:**
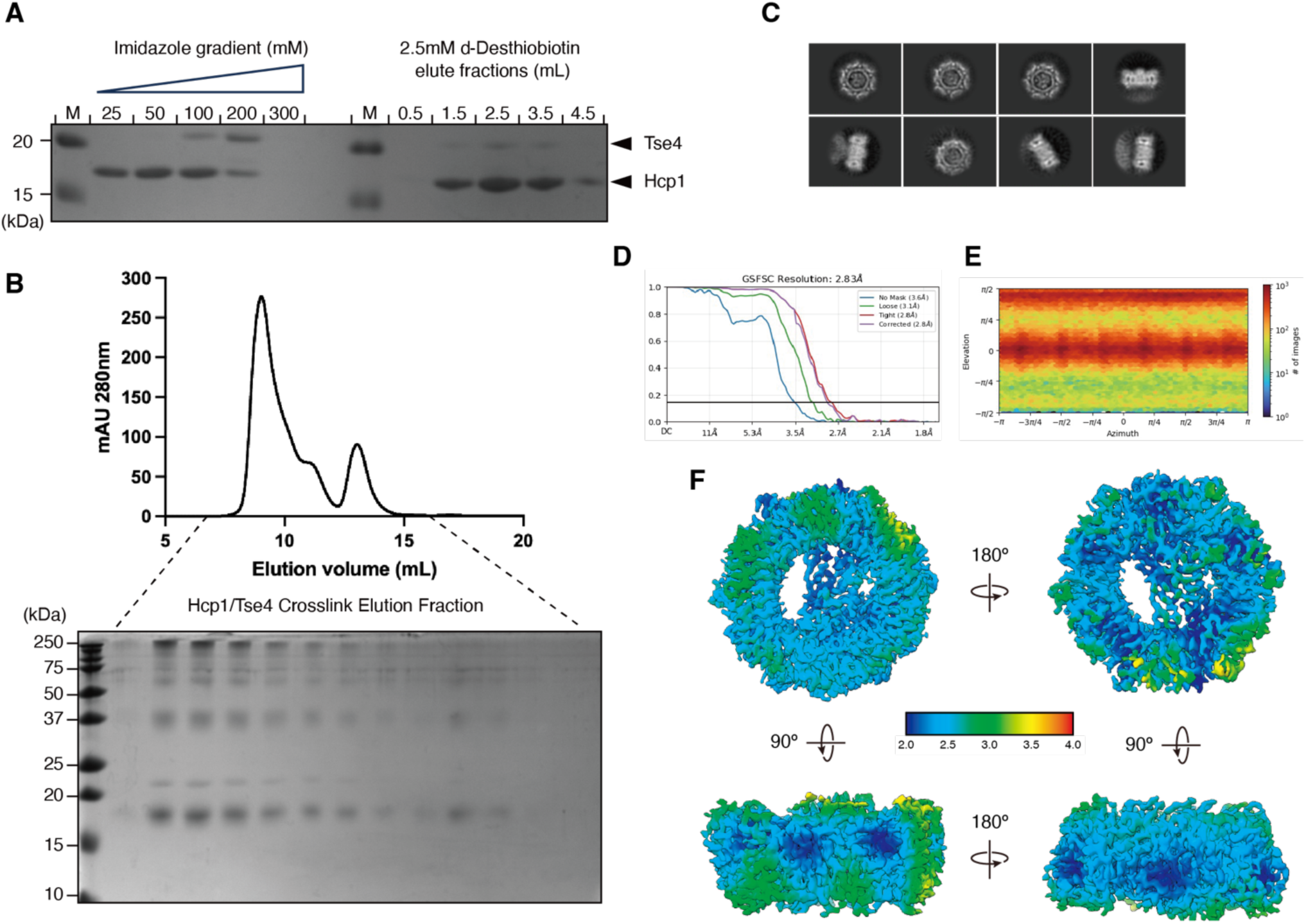
Biochemical purification, validation, and Cryo-EM characterization of the Hcp1-Tse4 complex. (A) SDS-PAGE analysis of Hcp1-Tse4 purification. Left panels show elution from Ni-NTA resin via an imidazole gradient (mM); right panels show elution from Strep-Tactin resin using d-Desthiobiotin (mL). Molecular weight markers (M) are indicated in kDa. (B) Size-exclusion chromatography (SEC) profile of the Hcp1-Tse4 complex. The elution volume is shown on the x-axis, with absorbance at 280 nm on the y-axis. SDS-PAGE analysis of the Hcp1/Tse4 crosslinked elution fractions corresponding to the SEC peaks, demonstrating the formation of high-molecular-weight assemblies. (C) Representative 2D class averages of Hcp1-Tse4. (D) Gold-standard FSC curve indicating a global resolution of 2.83 Å. (E) Euler angle distribution of the particles used in the final reconstruction. (F) Cryo-EM density map viewed from the top and side, colored by local resolution (Å).

**Extended Data Fig. 7:**
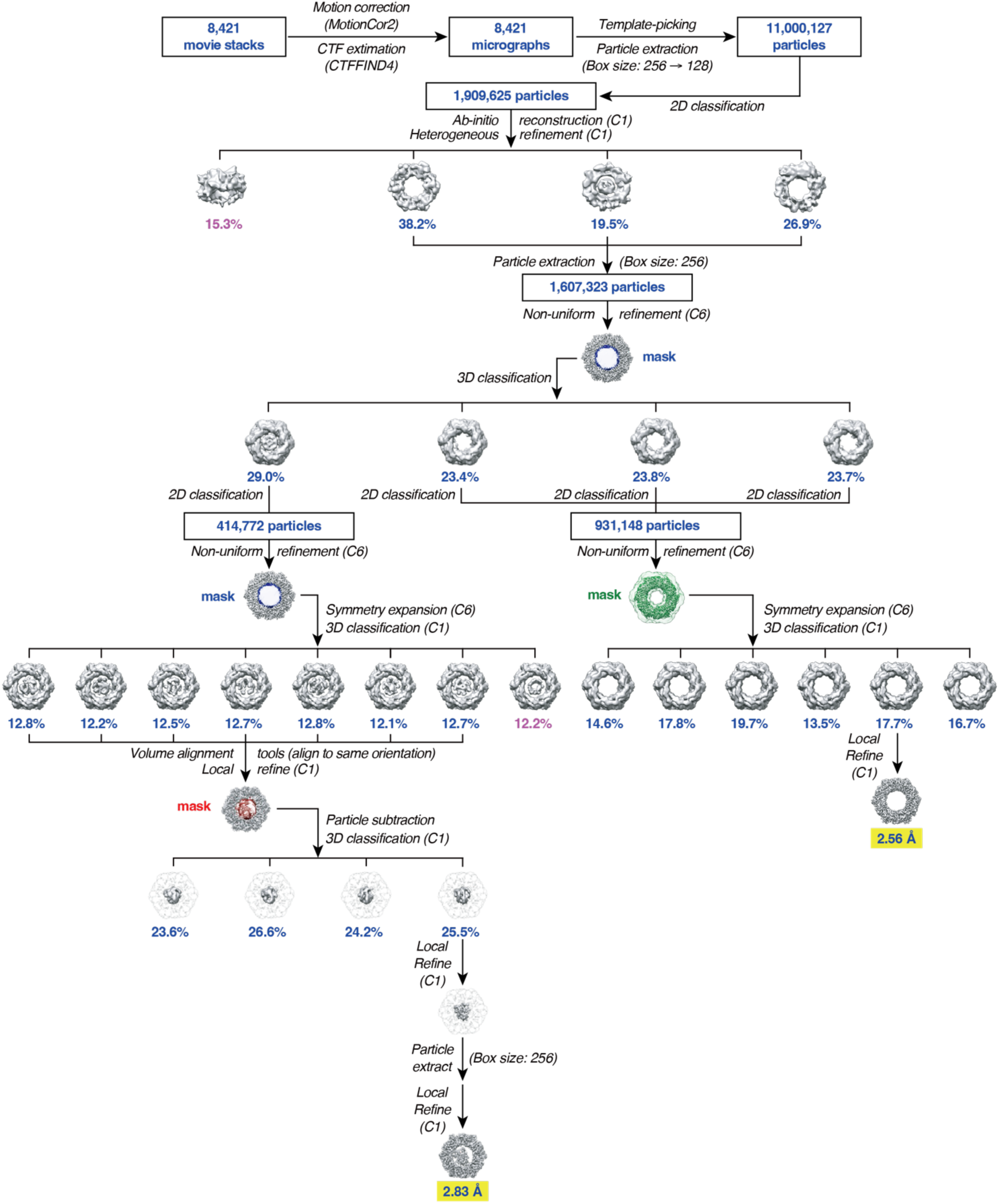
Cryo-EM data processing workflow for the Hcp1-Tse4 complex. Schematic representation of the Hcp1-Tse4 structural determination pipeline. The workflow outlines the progression from 8,421 movie stacks to the final high-resolution density maps. Following pre-processing, an initial 11,000,127 particles were refined down to 1,909,625 clean particles via iterative 2D classification and C1 heterogeneous refinement. A major class of 1,607,323 particles was selected for subsequent parallel refinement strategies: 1) Global Refinement: Symmetry expansion (C6) and focused 3D classification (masked in green) led to a masked local refinement (C1), producing a final 2.56 Å reconstruction. 2) Tse4-Specific Refinement: An alternative processing branch incorporated focused 3D classification, particle subtraction (masked in red), and multiple rounds of masked local refinement. Final particle extraction (256-pixel box) and focused refinement optimized the Tse4 density, resulting in a 2.83 Å reconstruction.

**Extended Data Fig. 8:**
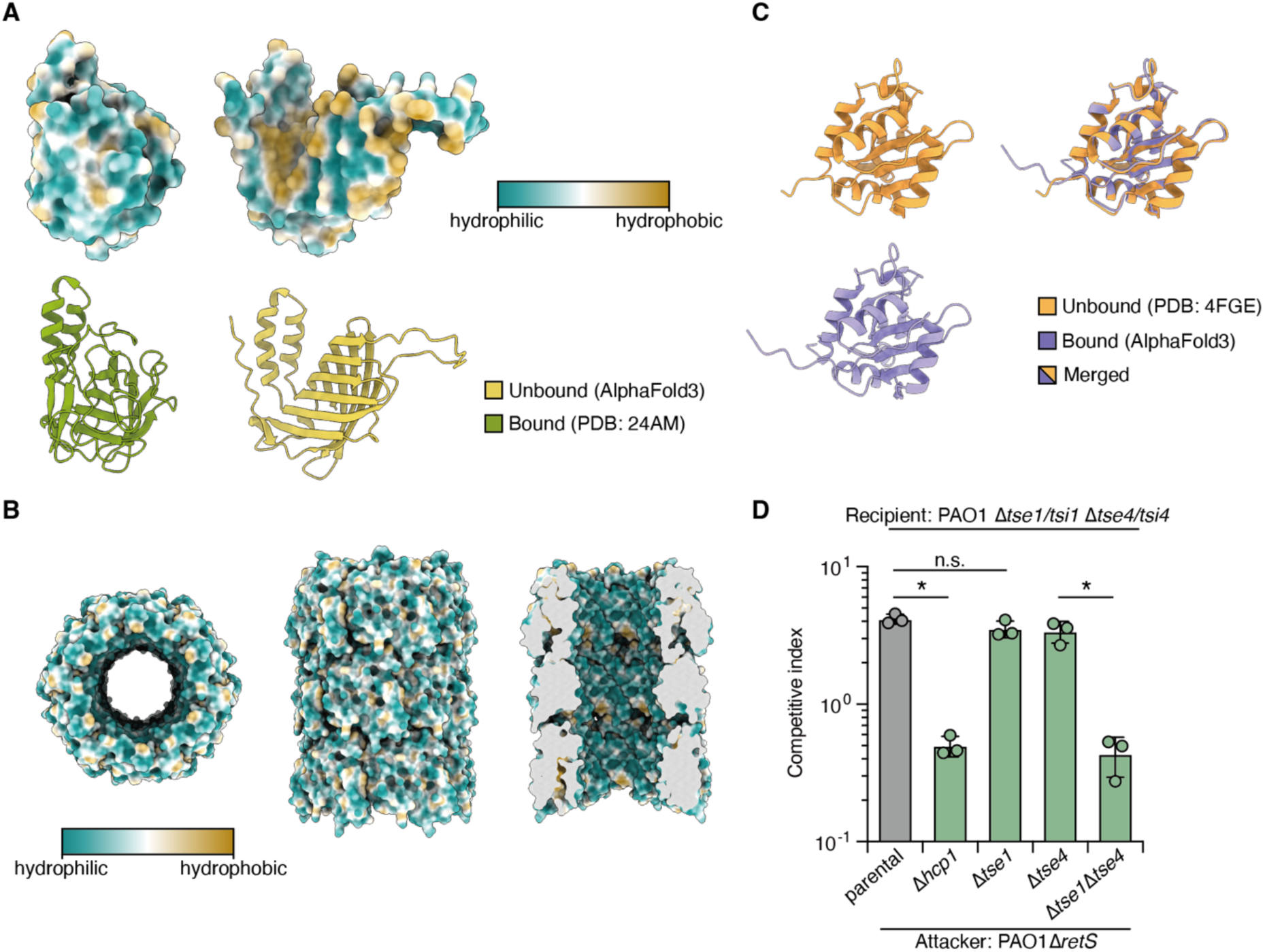
Structural analysis and functional validation of Hcp1-Tse1 and Hcp1-Tse4 complexes. (A) Surface hydrophobicity of Tse4 in Hcp1-unbound and -bound forms. Corresponding cartoons are shown below. (B) Surface hydrophobicity of Hcp1 inner-tube. (C) Tse1 in Hcp1-unbound and -bound forms. (D) Competitive index of PAO1 attacker strains. Cells were cocultured at a 10:1 attacker-to-recipient ratio for 6 hours. Data are presented as means ± SD. n = 3 biological replicates with three technical replicates each. Asterisks indicate statistically significant differences between the indicated mean values (p < 0.05, two-tailed t-test).

**Extended Data Fig. 9:**
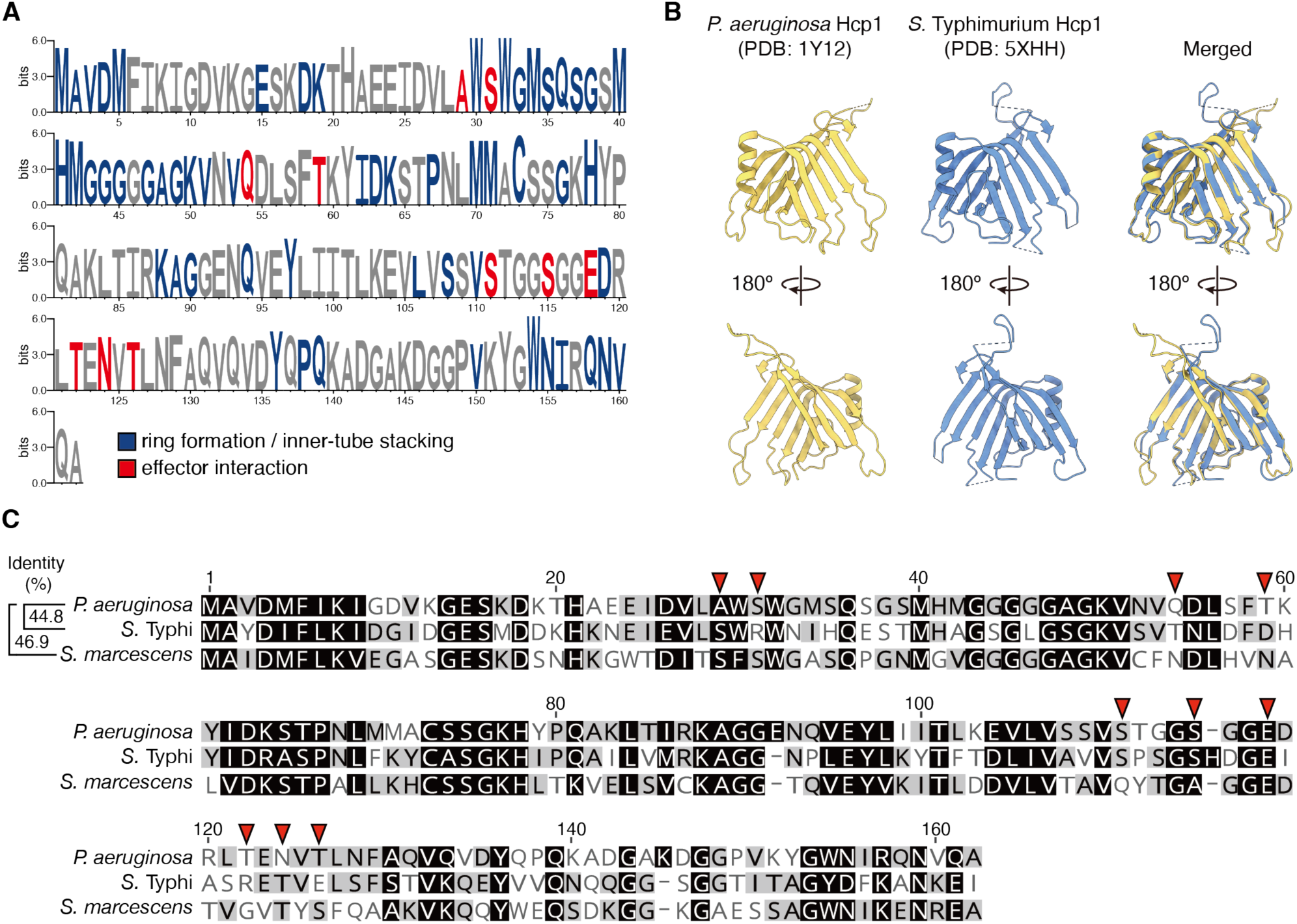
Sequence and structural comparison of Hcp1 homologs. (A) Sequence Logos of the Hcp1 multiple sequence alignments across homologs from *P. aeruginosa* isolates for 1,633 protein sequences, with residues involve in inner-tube structure assembly or effector interaction colored in blue and red, respectively. (B) Structural alignment of Hcp1 homologs from *P. aeruginosa* and *S.* Typhimurium. (C) Sequence alignment of Hcp1 homologs from *P. aeruginosa*, *S.* Typhi, and *S. marcescens.* The red arrows indicate the shared interface residues found within *P. aeruginosa* PAO1 Hcp1-effector complexes.

**Extended Data Fig. 10:**
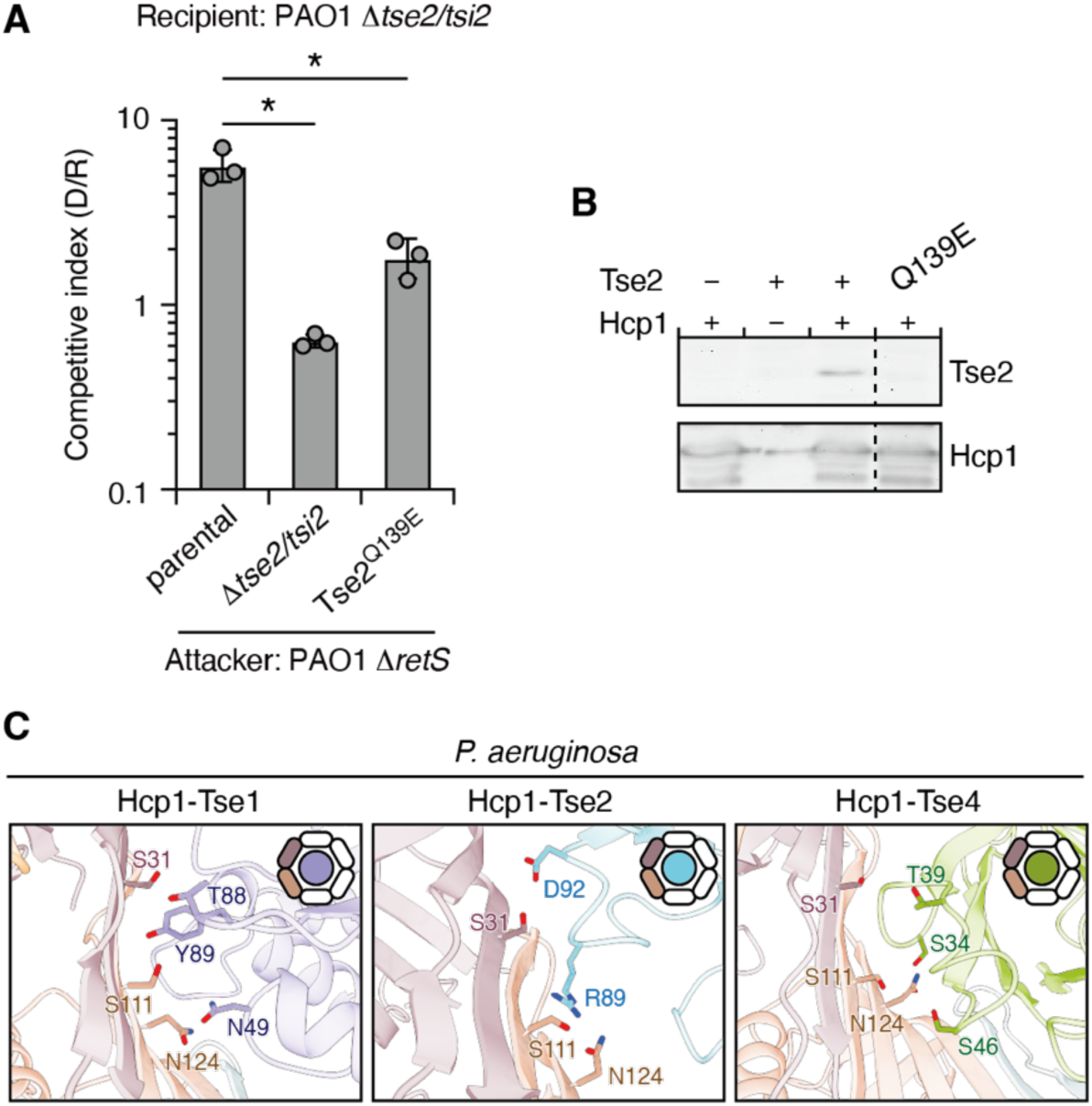
Shared effector-binding features are found across Hcp1 homologs. (A) Competitive index of PAO1 attacker strains. Cells were co-cultured with recipient cells at a 10:1 attacker-to-recipient ratio for 6 h. Data are presented as mean ± SD. n = 3 biological replicates, each with three technical replicates. Asterisks indicate statistically significant differences between the indicated groups (p < 0.05, two-tailed t-test). (B) Tse2 stability assay. A catalytic inactive Tse2 (T79A/S80A) was used as control. Tse2 or Tse2 carrying the Q139E substitution was co-expressed with Hcp1 in *E. coli*. Immunoblot analysis showing intracellular levels of Tse2 and Hcp1. (C) Conserved binding pocket across Hcp1–effector complexes from *P. aeruginosa*. Interface residues are labeled, and the corresponding Hcp1 subunits are highlighted in the ring schematics.

